# Fast bioluminescent nucleic acid detection using one-pot isothermal amplification and dCas9-based split luciferase complementation

**DOI:** 10.1101/2022.09.12.507659

**Authors:** Harmen J. van der Veer, Eva A. van Aalen, Claire M. S. Michielsen, Eva T. L. Hanckmann, Jeroen Deckers, Marcel M. G. J. van Borren, Jacky Flipse, Anne J. M. Loonen, Joost P. H. Schoeber, Maarten Merkx

**Affiliations:** Laboratory of Chemical Biology, Department of Biomedical Engineering, Eindhoven University of Technology, Eindhoven, The Netherlands; Institute for Complex Molecular Systems, Eindhoven University of Technology, Eindhoven, The Netherlands; Department of Clinical Chemistry, Rijnstate Hospital, Arnhem, The Netherlands; Laboratory for Medical Microbiology and Immunology, Rijnstate Hospital, Velp, The Netherlands; Research Group Applied Natural Sciences, Fontys University of Applied Sciences, Eindhoven, The Netherlands; Pathologie-DNA, Lab for Molecular Diagnostics, Location Jeroen Bosch Hospital, ’s-Hertogenbosch, The Netherlands

## Abstract

Nucleic acid detection methods based on isothermal amplification techniques show great potential for point-of-care diagnostic applications. However, most current methods rely on fluorescent or lateral flow assay readout, requiring external excitation or post-amplification reaction transfer. Here, we developed a bioluminescent nucleic acid sensor (LUNAS) platform in which target dsDNA is sequence-specifically detected by a pair of dCas9-based probes mediating split NanoLuc luciferase complementation. Whereas LUNAS itself features a detection limit of ∼1 pM for dsDNA targets, the LUNAS platform is easily integrated with recombinase polymerase amplification (RPA), providing attomolar sensitivity in a single-pot assay. We designed a one-pot RT-RPA-LUNAS assay for detecting SARS-CoV-2 RNA without the need for RNA isolation and demonstrated the diagnostic performance for COVID-19 patient nasopharyngeal swab samples using a digital camera to record the ratiometric signal. Detection of SARS-CoV-2 from samples with viral RNA loads of ∼200 cp/μL was achieved within ∼20 minutes, showing that RPA-LUNAS is attractive for point-of-care diagnostic applications.

---

**I**dentification of pathogens by detection of their nucleic acid fingerprints is a key strategy in clinical diagnostics, biomedical research, and food and environmental safety monitoring. The widely used quantitative polymerase chain reaction (qPCR) is highly sensitive, but requires expensive thermal cycling equipment and expert technicians, restricting its use to centralised laboratories that process samples sent in from local collection facilities. The resulting long time from sample to result and the limited diagnostic access in low-resource areas have stimulated the development of rapid, portable, and easy-to-use nucleic acid diagnostics that can be deployed at the point-of-care (POC)^1,2^.

Unlike PCR, isothermal nucleic acid amplification methods such as loop-mediated isothermal amplification (LAMP) and recombinase polymerase amplification (RPA) do not require thermal cycling for exponential accumulation of amplicons to detectable levels, and hence have gained interest for development of POC diagnostic tests^3–6^. Both LAMP and RPA are very rapid and highly sensitive techniques that operate at constant temperatures of ∼65°C and ∼40°C respectively. Although such low temperatures ease some requirements for equipment, they come with the risk of non-specific amplification, leading to potential spurious results when using nonspecific detection methods based on pH-change or fluorescent detection of total dsDNA. Therefore, various sequence-specific detection strategies have been developed for stringent target detection, including recent CRISPR diagnostic methods (CRISPR-Dx), which are mostly based on the collateral cleavage activity of type V (Cas12) and VI (Cas13) CRISPR effector proteins^7–10^. Following RPA- or LAMP-based target pre-amplification, Cas12 enzymes complexed with a short guide RNA (crRNA) bind sequence specifically to the amplicons, whereas Cas13 ribonucleoproteins (RNPs) require an additional in vitro transcription (IVT) step for converting dsDNA amplicons to target RNA^7,8^. Target binding in-turn triggers nonspecific nuclease activity, which is detected using cleavable fluorophore-quencher reporter nucleic acids^7,8,11^. However, the external excitation required for fluorescent detection gives rise to autofluorescence and scattering, limiting the sensitivity especially in complex media such as blood plasma^12^. Alternatively, the cleavage of reporter molecules can be visualised in a Laminar Flow Assay (LFA), which can be performed using simple paper-based devices and can be read by eye. However, LFA is inherently non-quantitative and prone to cross contamination due to post-amplification reaction transfer^11,13^.

Bioluminescent sensors require no external excitation and hence do not suffer from the sources of noise typical for fluorescence-based methods in complex samples. Bioluminescence also does not require sophisticated equipment but can be detected with any ordinary digital camera, making this type of readout particularly attractive for point-of-care applications^14–16^. Our group has previously reported the use of bioluminescence for detection of ssDNA and ssRNA targets through a luciferase conjugate molecular beacon, and a similar bioluminescent sensor concept based on toehold-mediated strand exchange was recently described by Winssinger and coworkers^17,18^. However, these methods are limited to detecting single stranded nucleic acids at picomolar levels, whereas attomolar sensitivity is typically required in diagnostics^19–21^. A bioluminescent dsDNA sensor based on dCas9-mediated split firefly luciferase complementation has been reported by Zhang and coworkers, but with a relatively modest limit of detection of ∼50 pM^22^.

Here we report LUNAS (Luminescent Nucleic Acid Sensor), a highly-improved generic platform for sequence specific, bioluminescent dsDNA detection based on dCas9-mediated split NanoLuc complementation, that can be easily combined with RPA isothermal amplification for one-pot real-time amplification and detection of nucleic acids (Fig. 1). In the RPA-LUNAS method, low input concentrations of target are efficiently amplified by (RT-)RPA, and the accumulating dsDNA amplicons serve as templates for recruitment of dCas9-LargeBiT and dCas9-SmallBiT ribonucleoproteins (RNPs) in close proximity, allowing reconstitution of the bright, blue light-emitting NanoLuc luciferase^23^. Addition of a green light-emitting calibrator luciferase previously developed in our group provides a robust blue-over-green ratiometric output^16^. RPA-LUNAS can be readily adapted to new targets, both DNA and RNA, by designing matching RPA primers and guide RNAs. Here, we demonstrate the performance of RT-RPA-LUNAS in SARS-CoV-2 detection, allowing direct detection of SARS-CoV-2 RNA in saliva, viral transport medium (VTM) and liquid amies medium (eSwab) without extraction. Finally, we demonstrate the detection of SARS-CoV-2 virus in COVID-19 patient samples within 10 – 30 minutes using an ordinary digital camera for readout, illustrating the potential for point-of-care application.

**Fig. 1:**
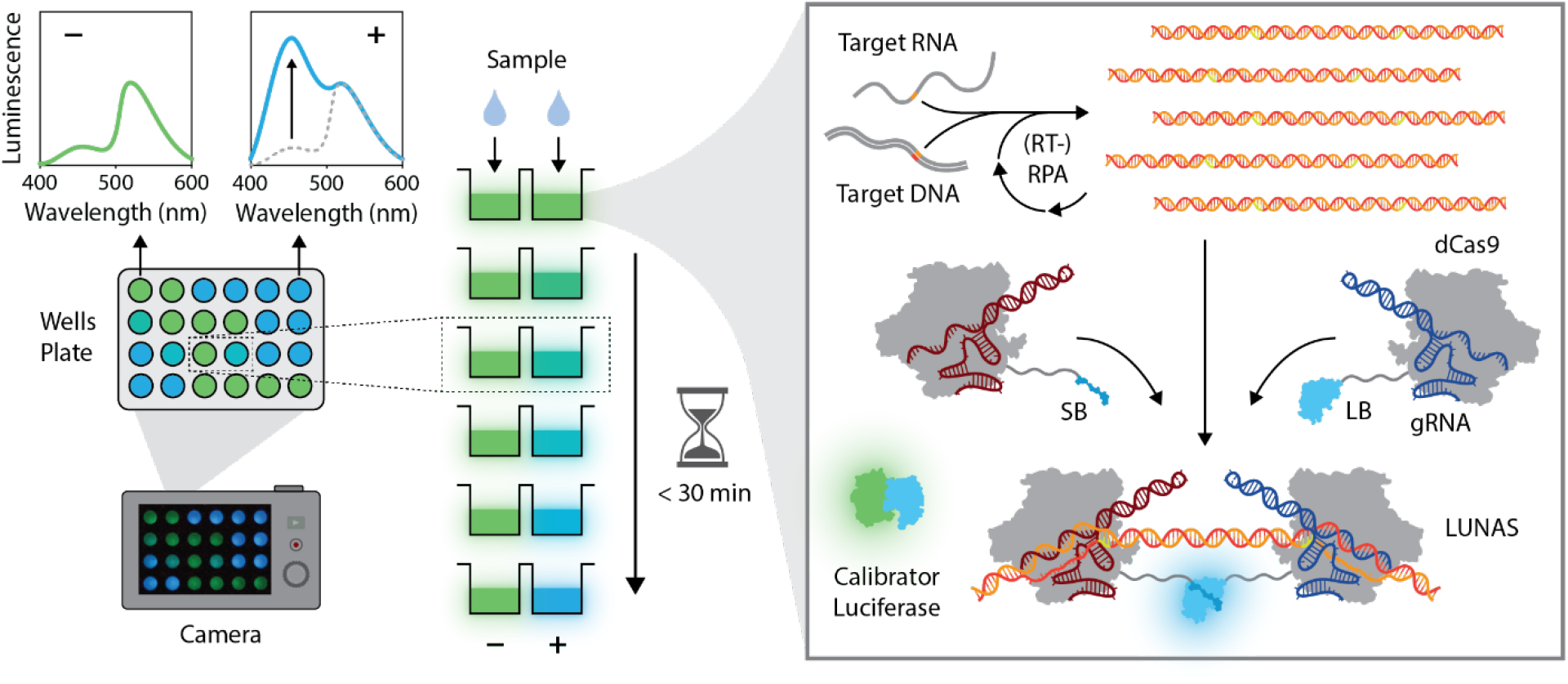
Schematic overview of the (RT-)RPA-LUNAS method. The presence of the target nucleic acid in a sample results in rapid exponential amplification of the target region through (RT-)RPA at a constant temperature of 40 °C. Two dCas9 proteins, fused to small BiT (SB) and large BiT (LB) split NanoLuc fragments respectively, and complexed with distinct guide RNAs (gRNA), bind to the resulting amplicons in close proximity of each other, allowing complementation of SB and LB and a resulting increase in blue luminescence (*λ*_*max*_∼ 460 nm) upon oxidation of the furimazine substrate. An mNeonGreen-NanoLuc (mNG-NL) fusion protein is included as a calibrator luciferase to correct for substrate turn-over, emitting green light (*λ*_*max*_∼ 520 nm) through Bioluminescence Resonance Energy Transfer (BRET). All components are combined in a one-pot reaction and, following the addition of a sample, the luminescence can be recorded by a digital camera. In the absence of target nucleic acid, no blue-light-emitting LUNAS complexes form, and mostly green luminescence is detected (low blue/green ratio), whereas the presence of the target nucleic acid results in an increase in blue signal (blue/green ratio high) within ∼ 30 min.

## Results

### LUNAS design and characterization

Prior to the advent of collateral cleavage based CRISPR diagnostics, Zhang and co-workers reported a bioluminescent dsDNA sensor based on dCas9-mediated split firefly luciferase complementation featuring a sensitivity of ∼50 pM^22^. We took this concept as a starting point for the design of a more sensitive sensor employing the brighter, smaller, and more stable split NanoLuc luciferase^19^. In our design, the 1.3 kDa small BiT (SB; *K*_*D*_= 2.5 μM) and 18.1 kDa large BiT (LB) luciferase fragments are fused to the C-terminus of *S. pyogenes* dCas9 via a flexible peptide linker (see Supplementary information for sequence). Complexation of the two dCas9 fusion proteins with distinct guide RNAs (gRNAs A & B) that are complementary to adjacent protospacer sequences in the target DNA enables the binding of the sensor RNPs to the DNA in close proximity of each other, resulting in the reconstitution of the luciferase. As gRNA exchange is known not to occur in vitro, separate incubation of dCas9-SB with gRNA_A and dCas9-LB with gRNA_B to form the sensor RNPs prior to the DNA sensing reaction avoids obtaining a statistical mixture of RNP variants wherein half of the resulting RNP:DNA complexes would not result in luciferase reconstitution^24^.

Expression of dCas9-LB and dCas9-SB was performed in *E. coli* and the proteins were purified by affinity column chromatography (Supplementary Fig. 3). To test the sensor and characterize its performance over a range of interspace distances between dCas9-SB and dCas9-LB, synthetic dsDNA fragments containing two protospacers on opposite strands spaced apart by 12 – 110 bp with PAMs facing inwards were used as target (Fig. 2A). We expected to attain the widest useful range of interspace distances using this protospacer orientation as the C-termini of the DNA-bound dCas9 proteins would face towards each other, minimising the distance to be bridged by the peptide linkers to allow for SB – LB complementation^25^. The dCas9-SB and dCas9-LB proteins were complexed with an excess of the complementary gRNAs (gRNA_T7A and gRNA_T7B, respectively) to form RNPs, and specific DNA binding was confirmed by an electrophoretic mobility shift assay (EMSA) (Supplementary Fig. 4). The RNPs (1 nM) were then incubated with target DNA (250 pM) for 30 minutes before the addition of furimazine substrate for signal detection. The sensor was found to yield maximal luminescent signal for interspace distances between 27 and 52 bp (Fig. 2B). Shorter distances presumably lead to steric hindrance between the two RNPs, whereas longer interspaces cannot be effectively bridged by the linkers to allow for split NanoLuc complementation, both resulting in low signal. Addition of control DNA lacking one or both of the protospacers did not result in any luciferase activation, confirming the sensor’s sequence specificity (Fig. 2B). The remaining background luminescence is likely due to low levels of split NanoLuc complementation driven by the intrinsic affinity of SB for LB (Supplementary Fig. 5).

**Fig. 2:**
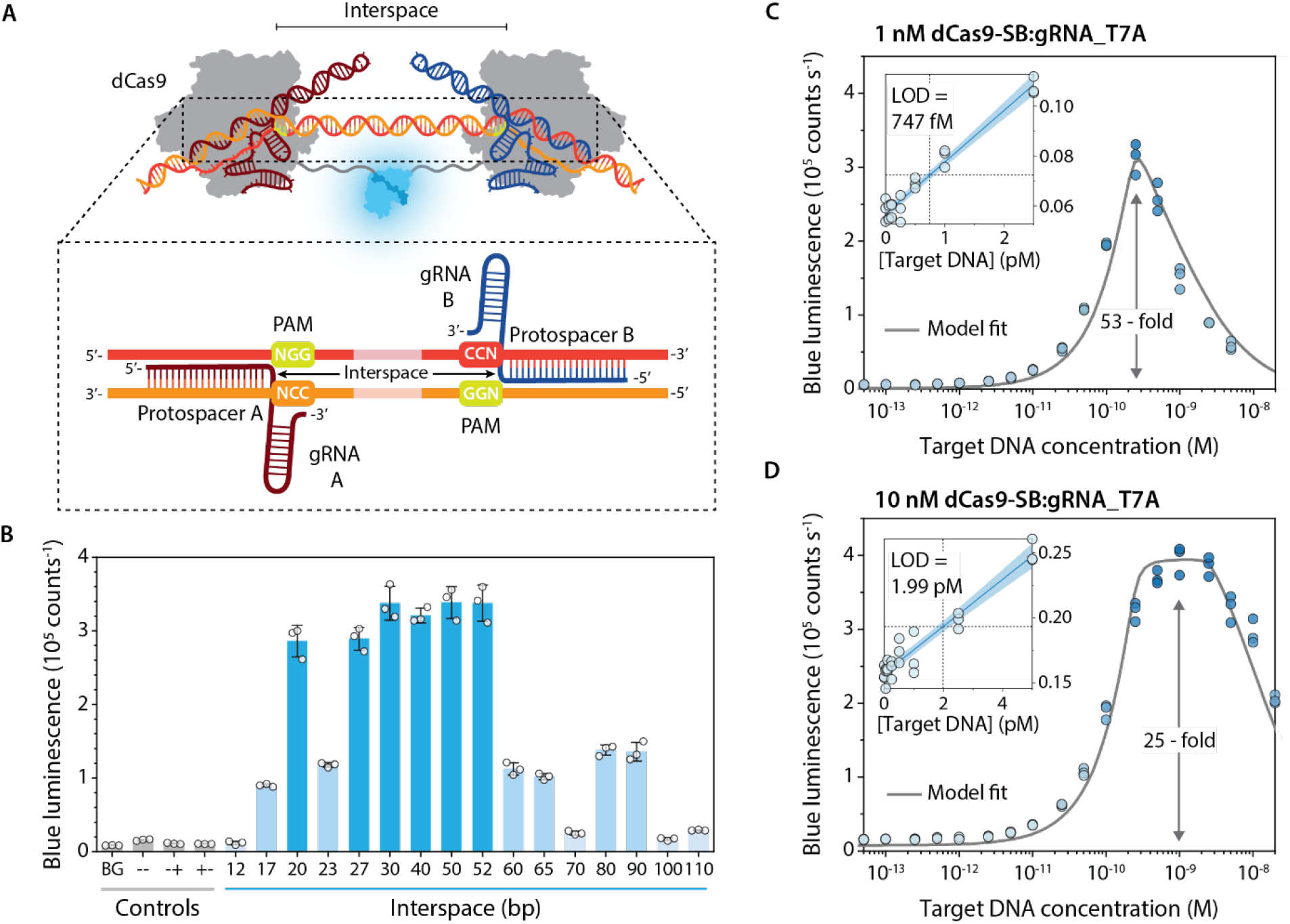
Characterisation of LUNAS. **A** Schematic overview of the LUNAS complex. The zoomed-in view of the dsDNA hybridized with the two gRNAs shows the orientation of the gRNAs with respect to each other on the target DNA and indicates the interspace region between the PAM-proximal ends of the gRNAs. **B** Bar chart showing the blue luminescence (458 nm) observed with 1 nM of both dCas9-SB and dCas9-LB sensor RNPs for targets with various interspace lengths and negative controls containing DNA with no target protospacer (−−), only one (−+) or the other (+−), or no DNA at all (background, BG). **C, D** LUNAS response curve for a target with an interspace of 50 bp, using **C** 1 nM dCas9-SB and dCas9-LB RNPs or **D** 10 nM dCas9-SB and 1 nM dCas9-LB RNP. A thermodynamic model was fitted to the data (grey solid line, see Supplementary Information). The limit of detection (LOD) was determined by local linear regression of sensor response at low target concentration (blue solid line with 95% confidence bands in inset). In (**B**), (**C**), (**D**), individual data points (*n* = 3, technical replicates) are represented as circles, and bars in (**B**) represent means, with error bars showing SD.

The concentration-response behaviour of the sensor was examined by incubating 1 nM sensor RNPs with a range of target DNA concentrations (50 bp interspace) for 30 minutes, to allow for sufficient time to reach equilibrium (Supplementary Fig. 6). A robust ∼50-fold maximum increase over background level was observed for 250 pM of target DNA (Fig. 2C). Higher DNA concentrations result in lower luminescence intensity, which is expected in case of analyte excess as the chance of both sensor RNPs binding to the same DNA molecule decreases with increasing excess of DNA^26^. Indeed, fitting a thermodynamic model of the system (see Supplementary information) yields a bell-shaped curve in close agreement with the data (Fig. 2C), assuming 25% of the RNPs is binding-competent, in accordance with previous reports on Sp(d)Cas9^27,28^. Model simulations also show this “hook” effect can be suppressed by increasing the relative amount of dCas9-SB RNP (Supplementary Fig. 2B), which we confirmed by measuring a concentration-response curve using 10 nM dCas9-SB and 1 nM dCas9-LB RNP (Figure 2D). The observed detection limit of 747 fM or 1.99 pM, using 1 or 10 nM dCas9-SB RNP respectively (Fig. 2C, D), represents a substantial improvement in sensitivity over prior bioluminescent nucleic acid sensors with LODs of ∼ 6 – 50 pM^17,18,22^. LUNAS is on par with fluorescent CRISPR Cas12- and Cas13-based methods excluding target pre-amplification, which feature LODs of ∼ 1 pM – 5 nM depending on the specific Cas12/Cas13 orthologs used, with recently reported sensitivities down to 166 fM only achieved by measuring for 1 – 2 hours^11,29,30^.

### Combining LUNAS with RPA

Although the ∼1 pM limit of detection of LUNAS is excellent for a direct nucleic acid sensor, the detection of pathogen nucleic acids from diagnostic samples typically requires sensitivity in the attomolar concentration range^19–21^. The sensitivity of LUNAS cannot be improved much further as it is limited by the minimal amount of reconstituted luciferase required for producing sufficient signal, which is on the order of 100 fM (Supplementary Fig. 7). Therefore, we explored combining LUNAS with isothermal amplification of the target nucleic acid. We selected RPA for its rapid exponential amplification at a relatively low optimal temperature of 37 – 42°C, a temperature at which both dCas9 and split NanoLuc are stable^23,31,32^. RPA primers were designed for one of the synthetic LUNAS target DNA fragments used in Fig. 2 (30 bp interspace). Initially, a two-step protocol was tested by first amplifying minute concentrations of target DNA during a 40-minute RPA reaction, followed by incubation of this reaction with the luminescent sensor mixture for 30 minutes at room temperature and subsequent measurement of luminescence intensity upon addition of substrate. This method provided excellent sensitivity, showing an increase in blue luminescence for inputs down to 2 copies of target DNA, detecting 3.3 aM in a 1 μL sample (Figure 3B). The assay response reached a plateau at an input ≥200 copies, which suggests that the RPA reaction was exhausted, yielding the same maximal amplicon output for higher input copy numbers^33^.

**Fig. 3:**
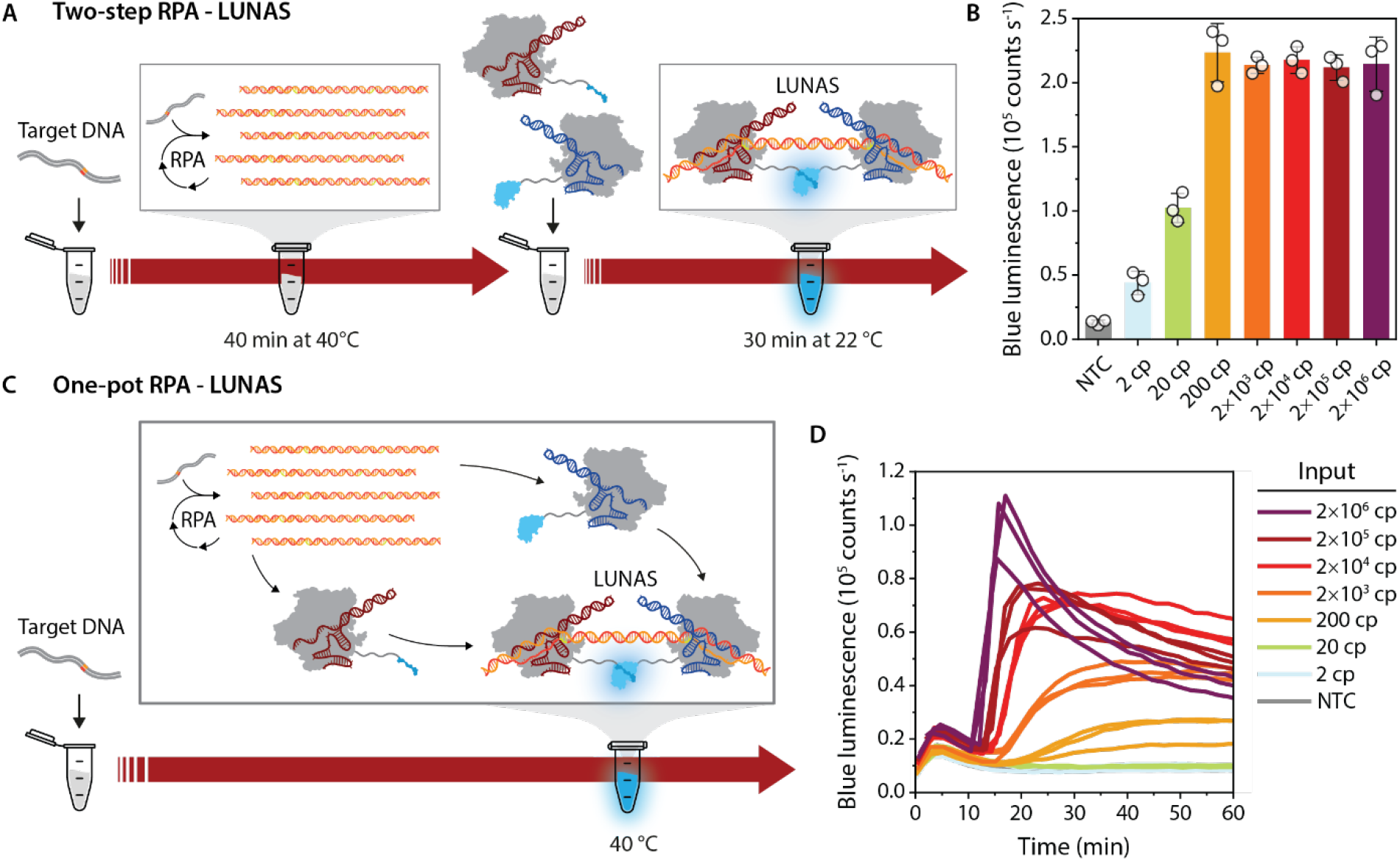
Combining LUNAS with RPA in two-step and one-step formats. **A** Schematic overview of the 2-step RPA – LUNAS method in which the target DNA is first amplified in an RPA reaction, which is subsequently added to a LUNAS reaction mixture for end-point detection of the amplicons. **B** Results of 2-step RPA – LUNAS assay response for a range of synthetic target DNA inputs (1 μL), using 10 nM dCas9-SB:gRNA_T7A and 1 nM dCas9-LB:gRNA_T7B in a 20 μL reaction. Individual data points (*n* = 3, technical replicates) are represented as circles, and bars represent means, with error bars showing SD. NTC: No template control. **C** Schematic overview of the one-pot RPA – LUNAS method in which both RPA and LUNAS components are present in the reaction mixture from the start, obviating post-amplification reaction transfer and allowing monitoring of the RPA reaction in real-time. **D** Results of one-pot RPA – LUNAS assay response over time for the same range of target DNA inputs as used in (**B**), using 10 nM dCas9-SB:gRNA_T7A and 1 nM dCas9-LB:gRNA_T7B. Individual replicate traces (*n* = 3) are shown.

Next we explored whether RPA and LUNAS amplicon detection could be performed simultaneously in a one-pot reaction. For this, the LUNAS reaction mixture including substrate was added to the RPA reaction, with RPA components at default concentrations, and luminescence was monitored while incubating the 20 μL mixture at 40 °C. Considering the multiple molecular interactions involved, with potential competition between binding of the dCas9 RNPs and components of the RPA reaction (recombinases, DNA polymerase, ssDNA binding proteins), the assay performed remarkably well. After an initial “bump” in luminescent signal, likely induced by the temperature increase in the wells, a sharp rise in signal was observed for high target input concentrations within 20 minutes, whereas target inputs down to 200 copies could still be clearly distinguished from the non-target control within 30 minutes (Fig. 3D).

### RT-RPA-LUNAS for SARS-CoV-2 RNA detection

Having achieved attomolar sensitivity in a one-pot method, we set out to design an RPA-LUNAS assay for the detection of a clinically relevant target. Triggered by the COVID-19 pandemic, we designed gRNAs and RPA primers for the SARS-CoV-2 RNA genome (Wuhan-Hu-1), taking into account prevalent nucleotide mutations known at the time and limiting homology with genomes of related human coronaviruses, including common cold viruses HCoV-OC43, HCoV-229E, HCoV-HKU1, and HCoV-NL63. Designed gRNA and primer sets targeting three different genomic regions were screened for the best performance using corresponding cDNA target fragments (Extended data Fig.1). The resulting best performing assay targets the ORF1a region, with primers flanking the sensor RNP binding sites (Fig. 4A). To allow for RNA detection, a reverse transcriptase was next included in the reaction mixture as well as RNase H, which degrades the RNA in DNA:RNA hybrids and was previously shown to enhance the sensitivity of RT-RPA reactions^34,35^. The resulting one-pot LUNAS assay was found to reliably detect as little as 200 copies of in vitro transcribed (IVT) ORF1a RNA fragment input in a 20 μL reaction within 25 minutes (figure 4B), with the two-step equivalent providing a further ∼10-fold increase in sensitivity (Extended Data Fig. 2). Lower inputs showed a stochastic assay response, with only some replicates yielding a detectable increase in signal.

**Fig. 4:**
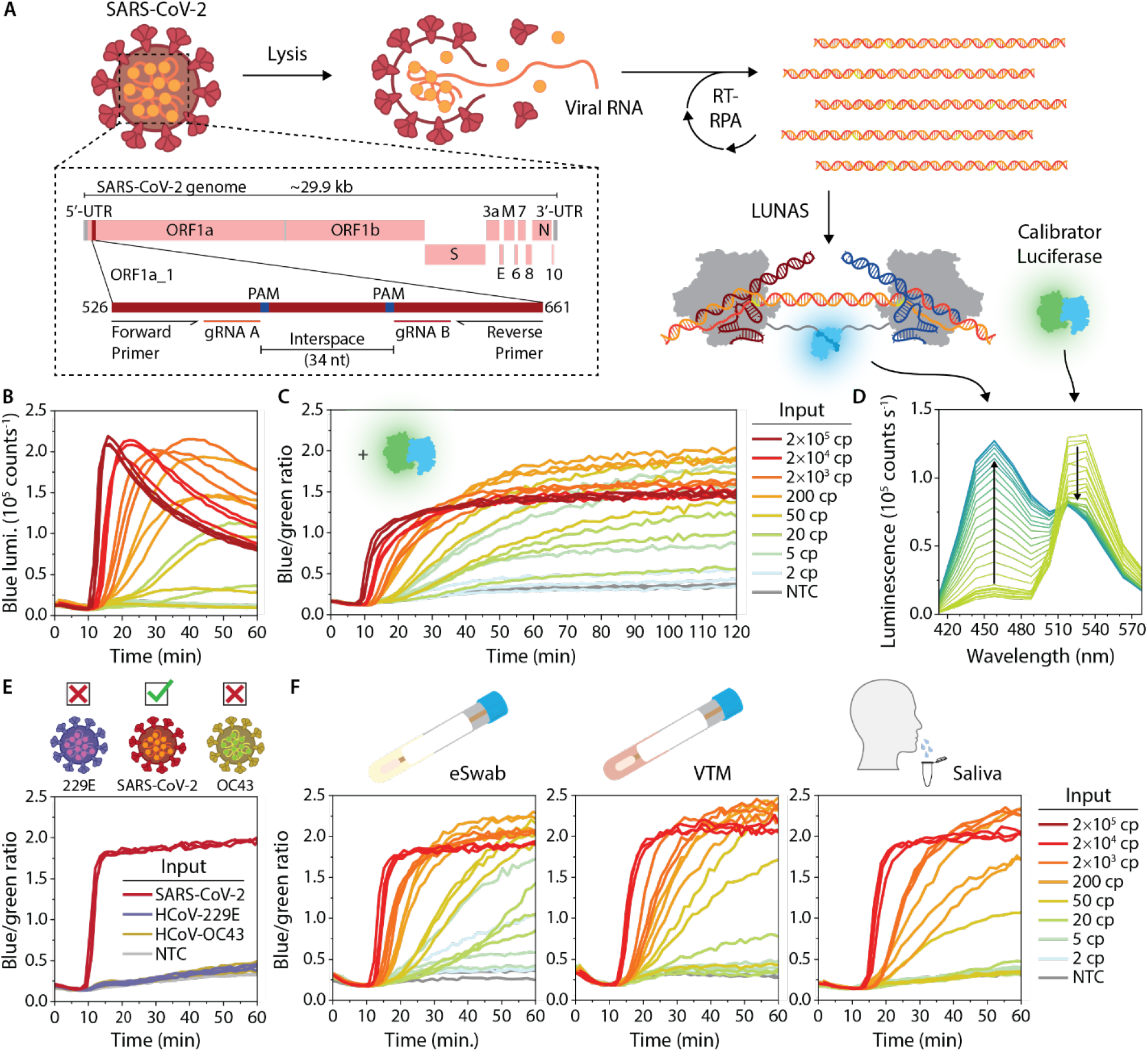
SARS-CoV-2 RT-RPA-LUNAS assay. **A** Schematic representation of the SARS-CoV-2 RT-RPA-LUNAS assay, showing the viral genome and zooming in on the assay target region (ORF1a_1) within ORF1a (see Supplementary Table 1 for primer and gRNA sequences). **B** Intensiometric one-pot RT-RPA-LUNAS response over time for a range of IVT ORF1a RNA inputs (see legend at **(C)**). **C** Ratiometric one-pot RT-RPA-LUNAS response over time for a range of IVT ORF1a RNA inputs. **D** Luminescence spectra of a single ratiometric one-pot RT-RPA-LUNAS reaction from panel (**C**) (200 cp input), showing one line per timepoint over the first 45 min, with trends indicated by arrows (see also Supplementary Fig. 8). **E** Ratiometric one-pot RT-RPA-LUNAS response over time for genomic RNA (2 × 10^4^ cp) isolated from SARS-CoV-2 (target) and from closely related HCoV-229E and HCoV-OC43 non-target viruses, showing the specificity for the target. **F** One-pot RT-RPA-LUNAS compatibility tests for eSwab, VTM and saliva sample matrices, showing ratiometric RT-RPA-LUNAS response over time for the same range of IVT ORF1a inputs as in (**D**). Mixtures of the sample media and inactivation buffer (1:1) were spiked with IVT target RNA and heated at 95°C for 5 min (eSwab, saliva) or 70°C for 10 min (VTM) (see methods) before addition to RT-RPA-LUNAS reactions. In (**B, C, E, F**) individual replicate traces (*n* = 3) are shown. NTC: No template control.

In the one-pot assay, the luminescence intensity gradually decreases over time after reaching a peak level, especially for the higher intensity signals. This time-dependence is non-ideal, in particular for point-of-care applications. First we considered the possibility that the decrease in signal was a reflection of the ‘hook’ effect observed in the titration experiment in Figure 2c, with amplicon accumulation leading to a decrease in signal upon redistribution of sensor RNPs over the larger number of amplicons. However, once bound to its target DNA, Sp(d)Cas9 is known to dissociate only very slowly, effectively staying tightly bound for hours^36,37^. We therefore performed an experiment in which the LUNAS sensor components were first incubated with an optimal concentration of target DNA (250 pM) for one hour. Subsequently, we increased the DNA target concentration further to 2.5 nM and continued monitoring the luminescence intensity. The rate of decrease in luminescence was not substantially faster compared to a control left at the initial target DNA concentration (Extended Data Fig. 3), indicating that binding of Sp(d)Cas9 RNP is indeed kinetically controlled. The decrease in luminescence intensity in RPA-LUNAS assays is therefore unlikely the result of redistribution of sensor RNPs over the larger number of amplicons. The decrease in absolute intensity in time may also be explained by gradual substrate depletion. Our group recently described the use of a green light emitting mNeonGreen-NanoLuc (mNG-NL) fusion protein, in which NanoLuc acts as a BRET donor to excite mNeonGreen, as a means for direct internal calibration of the RAPPID bioluminescent immunoassay^16,38^. Since the activity of this calibrator luciferase is equally dependent on the substrate concentration, including this calibrator luciferase in the RPA-LUNAS assay and taking the blue-over-green emission ratio yields an output corrected for substrate depletion. Indeed, doing so resulted in a ratiometric response that was stable in time, ruling out alternative explanations^39,40^ and simplifying result interpretation (Fig. 4C, D). Using this ratiometric assay, inputs as low as 5 cp could be detected, although stochasticity was observed for inputs < 50 cp (Fig. 4C).

To verify the specificity of the RT-RPA-LUNAS assay, the response for full genomic RNA isolates from SARS-CoV-2 as well as related HCoV-OC43 and HCoV-229E were compared in a one-pot experiment. Evidently, no increase in emission ratio was observed for all but the target input (Fig. 4E). With the aim to validate the assay using COVID-19 patient samples, we next explored suitable sample preparation methods. The typical RT-qPCR workflow involves chemical lysis and magnetic bead or filter based RNA isolation, which allows for control over the elution buffer and concentration of the RNA. However, in a POC setting a test should ideally require only a simple viral inactivation step to make the viral RNA accessible for detection. Successful heat-inactivation methods have previously been described for SARS-CoV-2 RNA detection from VTM (Viral Transport Medium), used for nasopharyngeal swab collection, and saliva^34,35^. In these methods, an inactivation buffer containing TCEP and an RNase inhibitor is used to effectively inactivate omnipresent RNases, notorious for being especially stable. To explore extraction-free detection from typical COVID-19 diagnostic sample matrices, we tested the compatibility of our assay with VTM, liquid Amies (eSwab) and saliva inputs. An inactivation buffer (100 mM TCEP, 1mM EDTA, 1U/μL murine RNase inhibitor, 10 mM Tris-HCl, pH 8.0) was added to the sample medium and IVT target RNA was added before heating the mixture to 95°C for 5 minutes (eSwab, saliva) or 70°C for 10 minutes (VTM). As a control, additional saliva samples were prepared without the addition of the inactivation buffer. For subsequent detection, 1 μL of a mock sample was used as input for a 20 μL RT-RPA-LUNAS reaction. In this way, target RNA could be detected from all three sample types, with the highest sensitivity for the eSwab sample, and approximately 10-fold lower sensitivity for saliva input (Fig. 4F). RNase inactivation is crucial in this procedure, as no target RNA was detected in saliva samples treated with heat only (Supplementary Fig. 9).

### SARS-CoV-2 assay validation

To clinically validate RT-RPA-LUNAS, we tested the diagnostic performance for nasopharyngeal patient samples. We used an experimental set-up using a heating block and a standard digital camera for signal detection, illustrating simple execution of the assay with minimal equipment (Fig. 5A). The camera-based readout was tested using the same conditions as for Fig. 4F, and similar responses were observed, with emission ratios calculated from blue- and green-channel intensities in RGB pictures (Fig. 5B and Supplementary Fig. 10). With this set-up an increase of blue signal was observed already after just 5 min for the higher concentrations, most likely reflecting the more efficient conductive heat transfer when using the heat block compared to the convective heating in the plate reader.

**Fig. 5:**
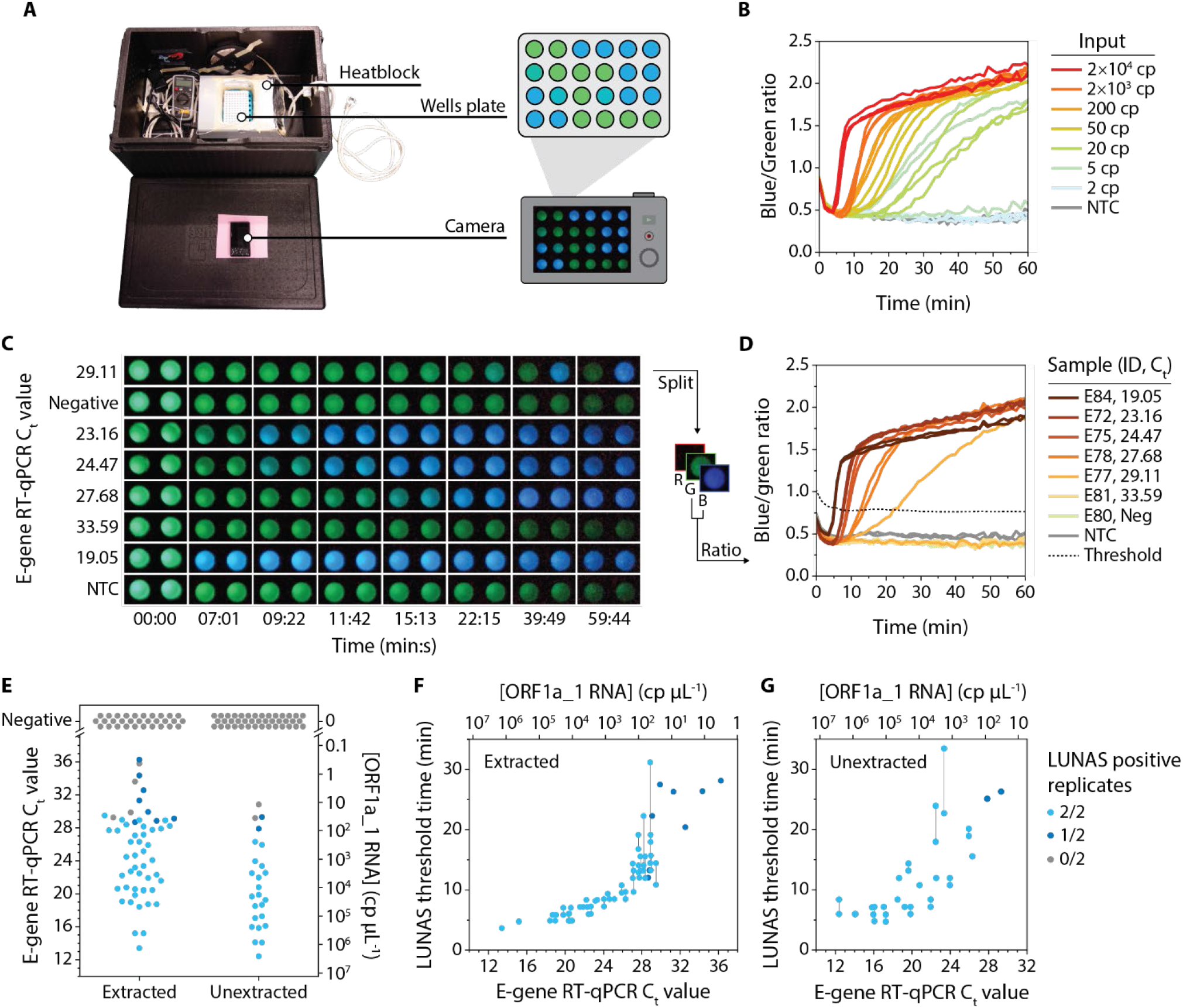
Detection of SARS-CoV-2 from patient samples. **A** Experimental set-up as used for camera-based readout of the ratiometric RT-RPA-LUNAS assay, consisting of a dark box with a simple heat block to keep a white 96-wells plate at 40 °C, and a digital camera fitted through a hole in the lid to continuously record pictures of the reactions. **B** Camera-based readout of experiment that is similar to the one shown in Fig. 4F, with ratiometric RT-RPA-LUNAS response extracted from pictures recorded by camera. Individual replicate traces (*n* = 3) are shown. **C** Pictures of a subset of ratiometric RT-RPA-LUNAS reactions with clinical sample inputs (RNA isolates), showing the blue/green intensity at several points in time. Two wells shown within the same dark box represent technical duplicates. **D** Blue/green ratio traces over time as extracted from the RGB images in (**C**). Individual traces are shown for both duplicates. The dashed line represents the threshold blue/green ratio and is based on the NTC and variance among all reactions (see methods). Also see timelapse in Supplementary Video. **E – G** Summary of RT-RPA-LUNAS test results from two independent groups of clinical samples: RNA extracted from nasopharyngeal swabs (‘extracted’) and eSwab samples that were treated according to our heat-inactivation protocol (i.e. ‘unextracted’). Colours code for RT-RPA-LUNAS outcome, and known E-gene RT-qPCR C_t_ values as well as the absolute concentration of the target region (ORF1a_1) in the original sample material predicted based on RT-ddPCR data for a subset of samples (Supplementary Fig. 11, Supplementary Table 2) are shown on the axes. **E** Overview of RT-RPA-LUNAS outcomes for all tested samples, showing a single data point per sample. **F** RT-RPA-LUNAS threshold time for all tested RNA isolates extracted from COVID-19 patient samples (see methods). **G** RT-RPA-LUNAS threshold time for all tested unextracted eSwab COVID-19 patient samples (see methods). In (**F, G**) datapoints represent individual replicates, with a vertical solid line connecting non-overlapping duplicates.

Using the camera-based readout, we next tested RT-RPA-LUNAS on clinical nasopharyngeal samples. Both RNA isolates (N=86, of which 55 positive), as well as eSwabs (N=66, of which 25 positive) treated by our heat-inactivation protocol directly prior to the assay, were included, constituting two independent sample groups for which comparator RT-qPCR results were known. To normalize and convert these C_t_ values to concentrations of the target RNA fragment, droplet digital PCR (RT-ddPCR) was performed for a selection of samples (Supplementary Fig. 11). Using the RT-RPA-LUNAS assay, the samples were measured in duplicate, adding only 1 μL per 20 μL reaction, and were considered positive for SARS-CoV-2 if the blue/green emission ratio surpassed the non-target control level by a threshold margin based on the variance among all reactions at t = 1 – 3 min (Fig. 5C, D) (see methods). All tested RNA isolates with an E-gene RT-qPCR C_t_ value < 28.5 (> 56 cp/μL; > 86 cp input) were identified as positive for SARS-CoV-2 by LUNAS within 22 minutes, with those having C_t_< 27 (> 154 cp/μL; > 231 cp input) detected within 11 minutes (Fig. 5E, F and Supplementary Fig. 12). The assay also performed well for the non-extracted samples, although based on the limited number of samples tested in the higher C_t_ value range the sensitivity appears to be slightly lower, correctly identifying all tested RT-qPCR-positive samples up to C_t_≈ 26.5 (> 215 cp/μL; > 107 cp input) (figure 5E, G, and Supplementary Fig. 13). We presume this difference in apparent limit of detection to be largely attributable to the 1.5-fold increase in RNA concentration occurring as part of the isolation procedure for the extracted samples versus the 2-fold dilution of samples in our heat inactivation protocol (see methods). For both types of input, all true negatives tested were indeed identified as such, including those positive for other respiratory viruses (see Supplementary Table 2). In samples for which the comparator assay reported a high C_t_ value, either one or both of the replicate reactions failed to signal the presence of SARS-CoV-2 RNA. Such stochasticity can also be observed for similar inputs (≤ 50 cp) of IVT target RNA (Fig. 4D, 4F, 5B), indicating similar sensitivity of the assay for the mock and the clinical samples. These results show that RT-RPA-LUNAS approaches the sensitivity of RT-qPCR, while providing important benefits with respect to assay time and required instrumentation, demonstrating its potential for point of care applications.

## Discussion

The LUNAS platform reported here provides a highly sensitive and generally applicable bioluminescent platform for sequence-specific dsDNA detection that is ideally suited to be used in combination with isothermal amplification methods such as RPA. RPA-LUNAS rivals other recently developed isothermal NA amplification- and CRISPR-based diagnostic methods in terms of speed, specificity and sensitivity, while providing in-sample bioluminescent detection that is particularly attractive for use in low resource settings. We showed that dCas9-split-NanoLuc based nucleic acid detection can be combined with RPA in an efficient one-pot assay with real-time readout, avoiding the risk of cross-contamination due to post-amplification reaction transfer. This is in contrast to lateral flow assay readouts or 2-step fluorescent approaches commonly used in isothermal amplification and CRISPR-based NA detection methods, which require transfer of the amplification reaction to the lateral flow device or the CRISPR cleavage reaction^34,35,41^. The readout of RPA-LUNAS is further simplified by including the previously developed mNeonGreen-NanoLuc calibrator luciferase^16,38^ to generate a robust ratiometric output, enabling straightforward signal recording by digital camera.

With two guide RNAs and two RPA primers that govern target recognition, covering a total of ∼100 nucleotides, RPA-LUNAS is highly specific, while also being easily programmable. Provided a 5’-*NGG-*3’ PAM and its reverse complement 5’-*CCN*-3’ exist within 27 – 52 nt of each other in a nucleic acid strand, this method can be readily adapted to detect such a target by designing corresponding gRNAs and RPA primers. As proof of principle, we demonstrated the use of RPA-LUNAS for the detection of SARS-CoV-2 RNA from nasopharyngeal samples of COVID-19 patients. Employing a simple heat-inactivation protocol, we showed that LUNAS is able to reliably identify COVID-19 positive cases without RNA isolation for samples with a viral load > 200 cp/μL, mostly within 20 minutes. This is on par with other CRISPR diagnostic methods applied to SARS-CoV-2 detection, such as the recent improved one-pot fluorescent SHERLOCK method (SHINE), which achieved similar sensitivity in 40 minutes (see Supplementary Table 3)^34^. For the specific case of the SARS-CoV-2 assay, the sensitivity can likely be improved further by increasing the sample input volume, decreasing the volume of the swab collection medium, or by higher-fold concentration in an RNA extraction step. Although progress has been made in simplifying RNA isolation in recent years, current methods are still cumbersome, leaving room for improvement for application in POC settings^42^.

The one-pot (RT-)RPA-LUNAS assay is remarkably effective considering the multiple potential interferences between reaction components, featuring a detection limit only ∼10-fold higher than the 2-step assay. This small difference in intrinsic sensitivity may be further reduced by additional optimization of the reaction conditions. The presence of dCas9 RNPs from the start of the amplification reaction could potentially block RPA for low target concentrations due to tight binding of the dCas9 complexes to the scarce templates. Such an effect has been observed for RPA / Cas12-based NA detection, and was recently resolved by introducing a photoactivatable guide RNA that allows for delayed activation of RNPs^43,44^. To enable multiplexing in a single reaction, a green or red color variant of the LUNAS system could be developed by use of BRET, analogous to the mNG-NL calibrator luciferase. Such a color variant could be used as part of a build-in control mechanism for appropriate sample collection to detect an endogenous NA that is amplified alongside the target NA in a duplex RPA reaction^46^

The simple assay set-up, with only a brief heat inactivation step for in-sample viral NA detection, is expected to allow for relatively straightforward integration of one-pot (RT-)RPA-LUNAS in a cheap and portable diagnostic device, that can be readout using a smartphone camera. Making use of advances in microfluidics and miniature electronic or chemical heating devices, integrated sample preprocessing, and lyophilization of the reaction mixture could further streamline the assay^46,47^. In conclusion, RPA-LUNAS shows great potential for translation into a sensitive and rapid point-of-care nucleic acid diagnostic that can readily be reconfigured for a quick diagnostic response following an epidemic outbreak.

## Methods

### Cloning and protein expression

The *S. pyogenes* dCas9 coding sequence was copied from the pET-dCas9-VP64-6xHis plasmid gifted by David Liu (Addgene plasmid #62935) by means of overhang PCR and was cloned via traditional restriction/ligation into a pET28a(+) plasmid (ordered from GenScript) coding for the C-terminal flexible linker and small BiT (SB) as well as large BiT (LB) and a C-terminal Strep-tag II (see protein coding sequence in Supplementary Information). *E*.*coli* BL21 (DE3) was transformed with the resulting plasmid directly for dCas9-SB expression, whereas the [SB – Strep-tag II – stop codon] sequence in-between the flexible linker and LB coding portions was removed by restriction/ligation for dCas9-LB expression. All cloning was confirmed successful by Sanger sequencing (BaseClear). Both proteins were expressed in *E*.*coli* BL21 (DE3), grown in LB-Miller medium with kanamycin (50 μg/mL) to OD_600_= 0.6 – 0.8 at 37°C and 160 rpm before induction by 0.2 M IPTG. Following overnight incubation at 18°C, 160 rpm, cells were harvested by centrifugation (8600×g, 15 min., 4°C) and resuspended in 12.5 mL pre-chilled lysis buffer (500 mM NaCl, 1 mM TCEP, 50 mM Tris-Cl pH 8.0) per gram of cell pellet, supplemented with benzonase (25 U/mL) (Merck) and a cOmplete EDTA-free protease inhibitor cocktail tablet (Merck). Cells were lysed by 3 passes through a high pressure homogenizer (Avestin Emulsiflex C3), at 15’000 – 20’000 Psi. The proteins were purified using Strep-Tactin XT (Iba) purification. Protein purity was confirmed by reducing SDS-PAGE (see SI) and concentrations were determined by measurement of absorbance at 280 nm on a NanoDrop spectrometer using extinction coefficients calculated from the protein sequence. Proteins in Strep-Tactin XT elution buffer (150 mM NaCl, 100 mM Tris-Cl pH 8.0, 1 mM EDTA, 50 mM D-biotin, 1 mM TCEP) were aliquoted and snap frozen in liquid nitrogen and stored at −70°C.

The mNG-NL calibrator luciferase was expressed and purified as described previously^16^, and the same procedure was followed for expression and purification of NanoLuc from a plasmid available in our lab (for Fig. S6).

### Assay design and synthetic target nucleic acids

Guide RNAs were designed using a custom Python tool based on the CRISPOR tool developed by Haeussler et al.^48^, taking into account the on-target activity as predicted by the scoring algorithms described by Moreno-Mateos et al.^49^ and Doench et al.^50^, as well as the specificity scores based on Hsu et al.^51^ and Doench et al.^50^. For the initial LUNAS assay used for sensor characterization, we designed 2 gRNAs that target bacteriophage T7 protospacers. These target sites were included in synthetic target DNA fragments with varying interspace distance separating the two protospacers. gRNA was generated from crRNA + tracrRNA (IDT) by combining both 1:1 in IDT nuclease free duplex buffer (30 mM HEPES, pH 7.5; 100 mM potassium acetate) to 4 μM final gRNA duplex concentration and heating at 95°C for 5 min., followed by gradually cooling to room temperature. gRNAs were aliquoted and stored at −30°C. Target DNA fragments were ordered as gBlocks (IDT) or PCR amplified from plasmids containing multiple such fragments, and then gel purified. For initial RPA-LUNAS experiments (Fig. 3), RPA primers were designed for these synthetic targets, following TwistAmp (TwistDx) primer design guidelines.

For the SARS-CoV-2 assay, the genome of the original Wuhan-Hu-1 isolate (GenBank: NC_045512.2) was scanned for suitable target site pairs using the guide RNA design tool. Candidate gRNAs were screened for specificity against genomes of related common cold human coronaviruses OC43, NL63, HKU1 and 229E as well as SARS-CoV and MERS-CoV (NCBI RefSeq accessions NC_006213.1; NC_005831.2; NC_006577.2; NC_002645.1; NC_019843.3; NC_004718.3 respectively). Three gRNA pairs predicted to be highly specific and having individual Moreno-Mateos^49^ activity scores >30/100 were selected. Complementary RPA primers were designed using the PrimedRPA tool developed by Higgins et al.^52^, aiming for small amplicon size. Final assay designs were checked for the absence of SNPs with >1% prevalence known at the time based on GISAID^53^/Nextstrain^54^ data in the UCSC SARS-CoV-2 genome browser^55^.

Corresponding SARS-CoV-2 cDNA fragments, PCR amplified from positive control plasmids provided by the FreeGenes project, were used for gRNA screening in LUNAS assays and subsequent primer screening in RPA-LUNAS assays. Synthetic ORF1a target RNA fragment was produced from the corresponding cDNA fragment by in vitro transcription using the HiScribe T7 High Yield RNA Synthesis Kit (NEB), according to manufacturer’s instructions. The IVT reaction was treated with DNAse I (Thermo Fisher) to degrade template DNA according to the manufacturer’s instructions. The IVT RNA was purified using a Monarch RNA cleanup spin column kit (NEB) and aliquoted for storage at −70°C. Concentrations of all nucleic acids were determined based on NanoDrop absorbance measurement at 260 nm (with 1 A260 optical density unit equal to 40 μg/mL RNA or 50 μg/mL dsDNA).

In addition, SARS-CoV-2 (Isolate USA-WA1/2020, BEI catalog No. NR-52347), HCoV-OC43 (BEI catalog No. NR-52727) and HCoV-229E (BEI catalog No. NR-52728) genomic RNA extracted from heat-inactivated virus was obtained from BEI Resources at known concentrations as determined by ddPCR.

The sequences of all crRNAs, RPA primers and synthetic targets are listed in Supplementary Table 1.

### Electrophoretic mobility shift assay (EMSA)

dCas9-SB:T7A and dCas9-LB:T7B RNP complexes were preassembled by incubating dCas9-SB with T7A gRNA and dCas9-LB with T7B gRNA separately in LUNAS buffer (20 mM Tris-Cl pH 7.5, 150 mM KCl, 5 mM MgCl_2_, 5% (v/v) glycerol, 1 mM DTT, 1 mg/mL BSA) for 15 min at 37°C, combining 1 μM protein with a 3-fold excess of gRNA. The EMSA was performed by incubating 12.5 nM target (473 bp) or non-target DNA (510 bp) fragment with 0.5 – 16 molar equivalents of dCas9-SB:T7A and/or dCas9-LB:T7B complexes in LUNAS buffer for 1 hour at room temperature. For subsequent electrophoresis, the reactions were supplemented with 1x loading dye (no SDS, NEB) and loaded onto a 2% agarose gel including 1x SYBR Safe, along with a 100 bp GeneRuler ladder (Thermo Fisher). Electrophoresis was performed for 1 h at 100 V in TAE running buffer (40 mM Tris, 20 mM acetic acid, 1 mM EDTA, pH 8.4). Bands were visualized on a Cytiva ImageQuant 800 using default settings for fluorescence imaging with automatic exposure adjustment.

### LUNAS assays

For LUNAS experiments, dCas9-SB:gRNA_A and dCas9-LB:gRNA_B complexes were preassembled separately using fresh protein and gRNA stock aliquots on the day of use by incubating dCas9-SB/LB (10 – 100x final sensor concentration) with the corresponding gRNA (3-fold excess) in LUNAS buffer (20 mM Tris-Cl pH 7.5, 150 mM KCl, 5 mM MgCl_2_, 5% (v/v) glycerol, 1 mM DTT, 1 mg/mL BSA) for 15 min at 37°C. LUNAS assays (without RPA) were performed at sensor RNP complex concentrations of 1 – 10 nM in a total volume of 20 μL, in Nunc 384-well non-treated flat-bottom white microplate (Thermo Fisher). Input DNA was prepared by serial dilution in LUNAS buffer, of which 1 μL was added to a LUNAS reaction. For the interspace variation and DNA titration assays (Fig. 2), 1 μL NanoGlo substrate (Promega, N1110) was added at a final dilution of 2100-fold after 30 min incubation at room temperature. Luminescence spectra (398 nm – 653 nm, step size 15 nm, bandwidth 25 nm) were recorded on a Tecan Spark 10M plate reader with an integration time of 100 – 200 ms and data was collected using Tecan SparkControl v2.1. ‘Blue’ luminescence refers to the luminescence intensity at 458 ± 12.5 nm. LODs were calculated in Microsoft Excel by linear regression of sensor response over a limited concentration range, using the standard deviation of the y-intercept^56^.

For the kinetic measurements (Fig. S5, S9), NanoGlo (1000-fold final dilution) was directly included upon mixing sensor RNP complexes and input DNA in a total reaction volume of 20 μL, and luminescence spectra were recorded over time.

### (RT-)RPA-LUNAS assays

For the 2-step (RT-)RPA-LUNAS assays, RPA reactions were prepared on ice using the TwistAmp Basic kit (TwistDx), first making a master mix comprising 505.26 nM of both primers, 14.74 nM magnesium acetate and 62.11% (v/v) TwistAmp rehydration buffer. For RT-RPA reactions, the master mix additionally included 2.11 U/μL SuperScript IV reverse transcriptase (Invitrogen), 0.105 U/μL RNase H (NEB) and 1.05 U/μL murine RNase inhibitor (NEB). This master mix was used to resuspend lyophilized RPA reaction components (TwistAmp Basic kit, TwistDx) at 47.5 μL per pellet. RPA reactions were performed at 40°C for 40 min in a total volume of 20 μL, combining 1 μL sample with 19 μL reaction mixture per replicate in a 96-well white PCR plate (VWR). For amplicon detection, a 1 μL sample was added to LUNAS reactions prepared and performed as described above.

For one-pot RPA-LUNAS assays, reactions were prepared on ice by first making a master mix containing 505.26 nM primers, 1053-fold diluted NanoGlo substrate, 14.74 nM magnesium acetate, 62.11% (v/v) TwistAmp rehydration buffer (TwistAmp Basic kit, TwistDx), and 10.53% (v/v) LUNAS mix (100 nM dCas9-SB:gRNA_A and 10 nM dCas9-LB:gRNA_B in LUNAS buffer). For ratiometric assays, 120 pM mNG-NL calibrator luciferase was included in the LUNAS mix. For RT-RPA-LUNAS reactions, the master mix additionally included 2.11 U/μL SuperScript IV reverse transcriptase (Invitrogen), 0.105 U/μL RNase H (NEB) and 1.05 U/μL murine RNase inhibitor (NEB). This master mix was used to resuspend lyophilized RPA reaction components (TwistAmp Basic kit, TwistDx) at 47.5 μL per pellet. Reactions were performed in a total volume of 20 μL, combining 1 μL sample with 19 μL reaction mixture in a Nunc 384-well non-treated flat-bottom white microplate (Thermo Fisher). Luminescence spectra (413 – 563 nm) were recorded over time at 40°C on a Tecan Spark 10M plate reader with an integration time of 100 ms. The blue/green emission ratio was calculated by dividing luminescence intensity at 458 ± 12.5 nm by the intensity at 518 ± 12.5 nm.

### Clinical validation

For testing clinical eSwab (Copan, Italy) samples without RNA isolation, viral lysis and nuclease inactivation was performed by adding an inactivation buffer (200 mM TCEP, 2 mM EDTA, 2 U/μL murine RNase inhibitor, 20 mM Tris-HCl, pH 8.0) to the sample in 1:1 ratio, followed by incubation at 95°C for 5 min. The compatibility of RT-RPA-LUNAS with this sample pretreatment method, based on Arizti-Sanz et al.^34^ and Qian et al.^35^, was tested using mock eSwab, as well as VTM (HiMedia HiViral medium) and saliva (obtained from a healthy donor) samples, which were prepared by spiking in IVT ORF1a target RNA. Target RNA was added before heating in order to simulate release of RNA from lysed virus before complete denaturation of RNases.

Clinical RNA isolate samples were extracted from nasopharyngeal swabs collected in either 1 mL eSwab or 2 mL PurePrep TL+ buffer (MolGen). For eSwabs, 150 μL sample was combined with 150 μL MP96 external lysis buffer (Roche) for viral lysis. RNA was extracted from 300 μL of this mixture or from 300 μL of samples collected in PurePrep TL+ directly using the STARMag 96×4 Viral DNA/RNA Universal kit (SeeGene, South-Korea) on a Hamilton Starlet, and was eluted in 100 μL.

The one-pot ratiometric RT-RPA-LUNAS assay was prepared as described above, with a higher calibrator luciferase (24 pM final concentration) and NanoGlo substrate concentration (400-fold final dilution). One or two NTC reactions (water input) were included per assay run of 18 – 56 duplicate reactions. Samples (1 μL) were added to reaction mixtures (19 μL) in a 96-well white PCR plate (VWR) on ice, which was then briefly centrifuged in a lettuce spinner^57^ before placing it in a heating block set at 40°C (monitored by external thermometer) within a black EPP box to exclude ambient light. Luminescence was recorded using a Sony DSC-RX100 III digital camera fitted through a hole in the lid of the box, with 30s exposure time, f/1.8 and ISO-6400. The camera was controlled by the Sony Imaging Edge Mobile app on an Android device with an auto clicker app (Click Assistant - Auto Clicker by Y.C. Studio) for continuous shooting over 1 hour. Resulting RAW images were converted to 16 bit TIFF files using Sony Imaging Edge Desktop, and mean blue (B) and green (G) intensities per well were extracted from split RGB channels in ImageJ (v1.53q). Reactions were regarded as positive for SARS-CoV-2 if the blue/green ratio (BG(t_i_)) exceeded a threshold value BG_T_(t_i_) = BG_smoothNTC-_(t_i_) + 6×SD_All,1-3min_ for t_i_ to t_i+2_, with BG_smoothNTC_(t_i_) the moving average blue/green ratio of the NTC over (t_i-5_, t_i+5_), and SD_All,1-3min_ the standard deviation of the blue/green ratio of all reactions between t = 1 min and t = 3 min. The reported LUNAS threshold time t_T_ equals the first t_i_ satisfying the condition BG(t_i_) > BG_T_(t_i_) for (t_i_, t_i+2_).

### RT-qPCR and ddPCR testing of clinical samples

RNA was extracted as described above, followed by RT-PCR using the Allplex SARS-CoV-2 assay (SeeGene, South-Korea) on a CFX96 thermocycler (Biorad), simultaneously detecting four different target genes: E, N, RdRp and S (the latter two are combined in one fluorescent signal). Data was analyzed with the SARS-CoV-2 Viewer software (SeeGene). Some samples were initially tested using the BioFire Respiratory Panel 2.1+ (bioMérieux, France), enabling qualitative detection of multiple pathogens.

To quantify absolute ORF1a_1 RT-RPA-LUNAS target concentrations and correlate these with RT-qPCR C_t_ values, droplet-digital PCR was performed for 16 extracted RNA samples using the 1-Step RT-ddPCR Advanced Kit for Probes (BioRad) and the CFX96 thermocycler (BioRad) and QX200 ddPCR system (BioRad). Three different reactions were performed per sample, targeting the E- and the N-gene as well as the ORF1a_1 region targeted by the RT-RPA-LUNAS assay. Samples were diluted 1 – 1000-fold to avoid overloading and 5 μL of diluted sample was added per 22 μL PCR reaction. Data was analysed using BioRad QX One (v1.2) and linear regression of the correlation between resulting concentrations and RT-qPCR C_t_ values was performed in Microsoft Excel.

All results are listed in Supplementary Table 2.

### Ethics statement

The use of anonymised clinical samples for this study was evaluated and approved by the local medical ethics review committee of the Rijnstate Hospital (reference number: KCHL 2021-1950), and samples were acquired in accordance with the Declaration of Helsinki. The samples used here were obtained as part of standard clinical COVID-19 testing and patients did not object to the use of remnant sample material for quality purposes and research.

### Material availability

Plasmids for recombinant expression of dCas9-SB and dCas9-LB will be made available through AddGene after publication. Protein and DNA sequences are available in the Supplementary Information.

## Data availability

Source data are provided with this paper.

## Code availability

The adaptation of the CRISPOR^48^ command line gRNA design tool used in this study will be made available on GitHub after publication.

## Acknowledgements

We especially thank the other members of the iGEM 2019 Eindhoven University of Technology team Jo-Anne Ewald, Noëlle Gerards, Anouk Marinus, Roy van Mierlo, Yvonne van Mil, Mandy Shao, and Chris Tomassen for their contribution to the iGEM project that lies at the base of this work. We thank Juliëtte Schaghecke (Fontys University of Applied Sciences, Eindhoven, the Netherlands) for her help in performing RT-ddPCR. We are thankful to the FreeGenes project for providing SARS-CoV-2 positive control plasmids. The following reagents were obtained through BEI Resources, NIAID, NIH: Genomic RNA from HCoV-OC43 (NR-52727), HCoV-229E (NR-52728), and SARS-CoV-2 (NR-52347). This work was supported by RAAK.PRO Printing makes sense (RAAK.PRO02.066) and funding from the Eindhoven University Fund (COVID19 Engineering Fund).

## Author contributions

H.v.d.V. conceived and designed the study, performed experiments, analyzed the data, and wrote the manuscript. E.v.A. supervised early experiments for this work that provided proof of principle and contributed to the clinical validation experiments. C.M., E.H., and J.D. contributed to the design of LUNAS and initial LUNAS experiments. M.v.B. and J.F. supervised clinical validation work, and J.F. provided RT-qPCR data and important feedback. A.N. and J.S. supervised initial SARS-CoV-2 assay experiments and provided important feedback. M.M. conceived, designed, and supervised the study, analyzed the data, and wrote the manuscript. All authors discussed the results and commented on the manuscript.

## Extended data

**Extended Data Fig. 1:**
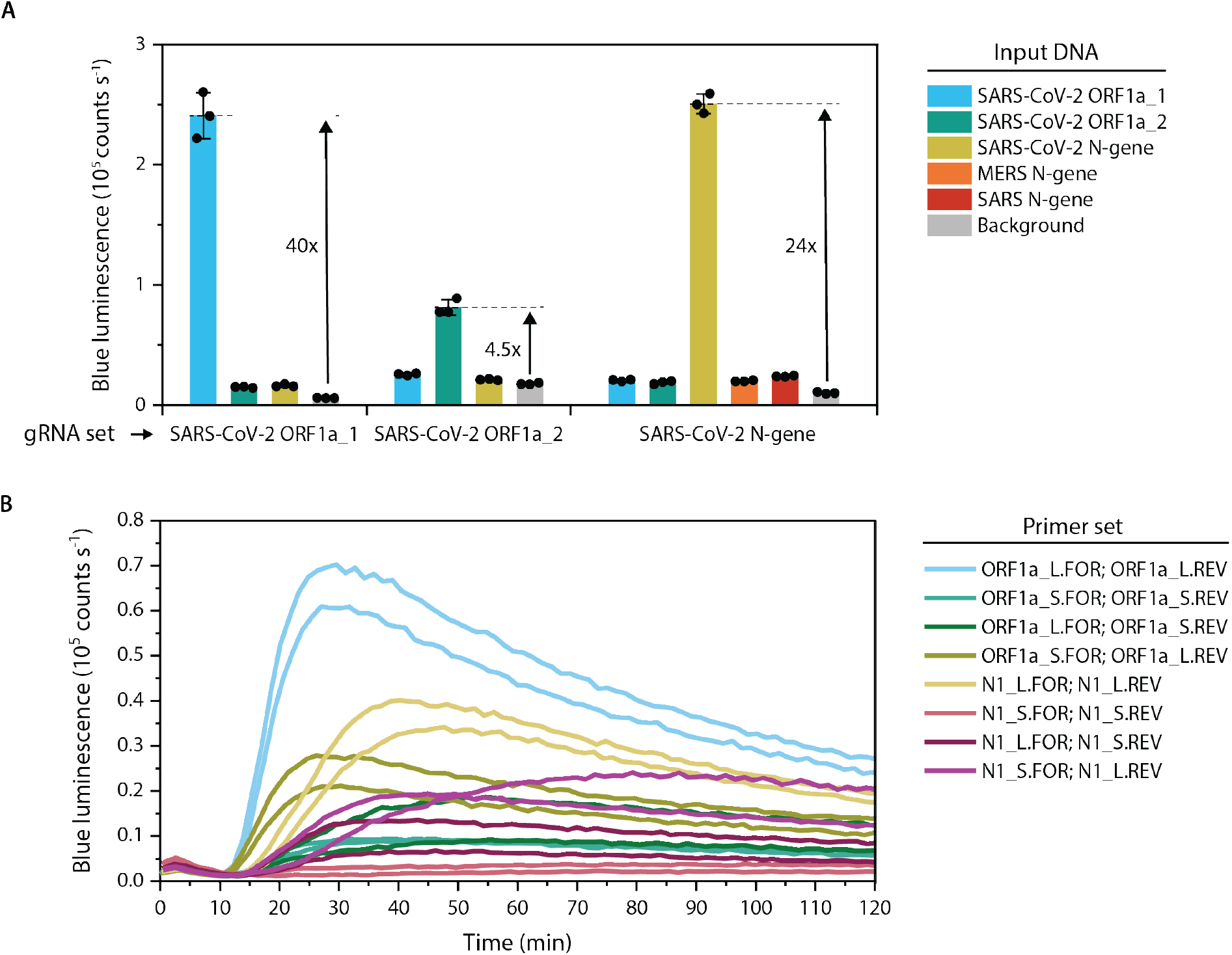
RPA-LUNAS SARS-CoV-2 assay gRNA and primer screening. **A** Screening 3 pairs of gRNAs for best performance in a LUNAS assay. dCas9-LB and dCas9-SB proteins were complexed with 3 pairs of gRNAs targeting different regions in the SARS-CoV-2 genome (ORF1a_1 and ORF1a_2 designate two different regions of ORF1a, see Supplementary Table 1). With these dCas9 RNPs (1 nM of both SB and LB), LUNAS assays were performed on matching target cDNA fragments (250 pM), as well as on the non-matching fragments (250 pM) used as non-target controls. Additionally, for the N-gene LUNAS assay, MERS and SARS N-gene cDNA fragments (250 pM) were used as non-target controls featuring strong sequence-homology with the target. Clearly, the ORF1a_1 and N-gene gRNA sets show the best LUNAS performance, with a 40- and 24-fold increase in blue luminescence over background level respectively, while the ORF1a_2 set only shows a moderate 4.5-fold increase and low absolute intensity. The low response of the SARS-CoV-2 N-gene LUNAS for MERS and SARS N-gene cDNA fragments compared to the target SARS-CoV-2 N-gene cDNA fragment demonstrates the assay specificity. Bars represent means of technical replicates (n = 3), which are indicated as black dots. Error bars show SD. **B** Screening RPA primer sets for best performance in an RPA-LUNAS assay. Two forward and reverse primers were designed for combination with the two best performing SARS-CoV-2 LUNAS assays in (A), one partly complementary to the PAM-distal protospacer sequence (‘_S’ primers), the other only binding to the sequence 3’ from the protospacer (‘_L’ primers). All 4 combinations of primers per target region were tested. For both the N and ORF1a target region, the assays using the combination of only ‘_L’ primers showed the highest signal. Clearly, the ORF1a assay performed best, showing the quickest onset of signal increase, and reaching the highest absolute intensity. The assays were performed at 42°C, using 200 cp of input cDNA fragment. Individual replicate traces are shown (n = 2).

**Extended Data Fig. 2:**
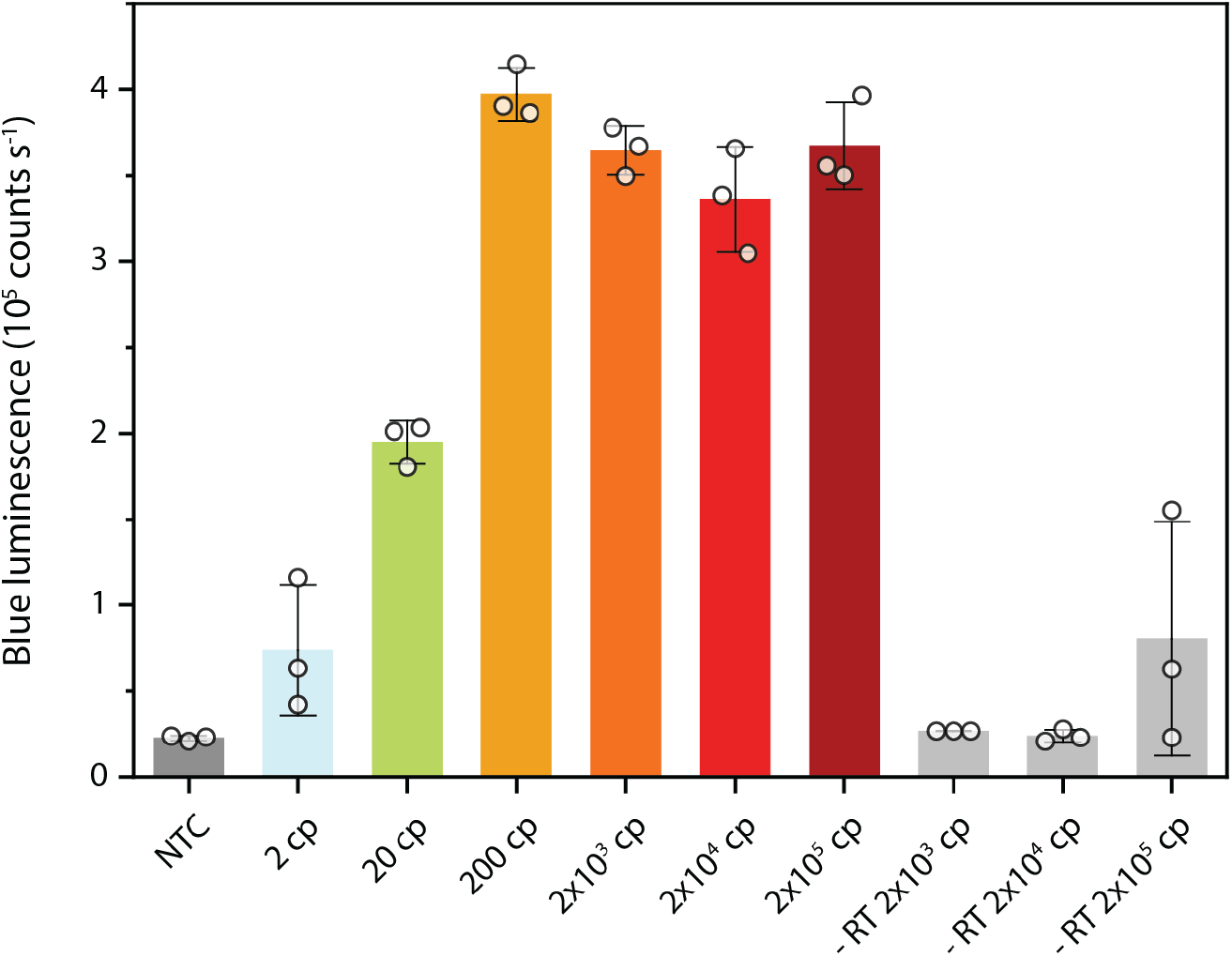
Two-step RT-RPA-LUNAS SARS-CoV-2 assay. RT-RPA reactions were performed for a range of inputs of IVT SARS-CoV-2 ORF1a RNA fragment. Additionally, control reactions in the absence of the reverse transcriptase (labelled ‘-RT’) were performed to verify successful DNase I-based degradation of IVT template DNA. The triplicate RPA reactions were added to LUNAS assay reactions and resulting blue luminescence was measured. For the reactions without RT, only for an input of 2×10^5^ cp a LUNAS response is observed for 2 of 3 replicates, similar to the response for 2 cp input in reactions including RT, confirming that the RT-RPA-LUNAS assay indeed detects RNA and that there is virtually no contribution of remainder IVT template DNA to the observed results. Bars represent means of technical replicates (n = 3), which are indicated as circles. Error bars show SD.

**Extended Data Fig. 3:**
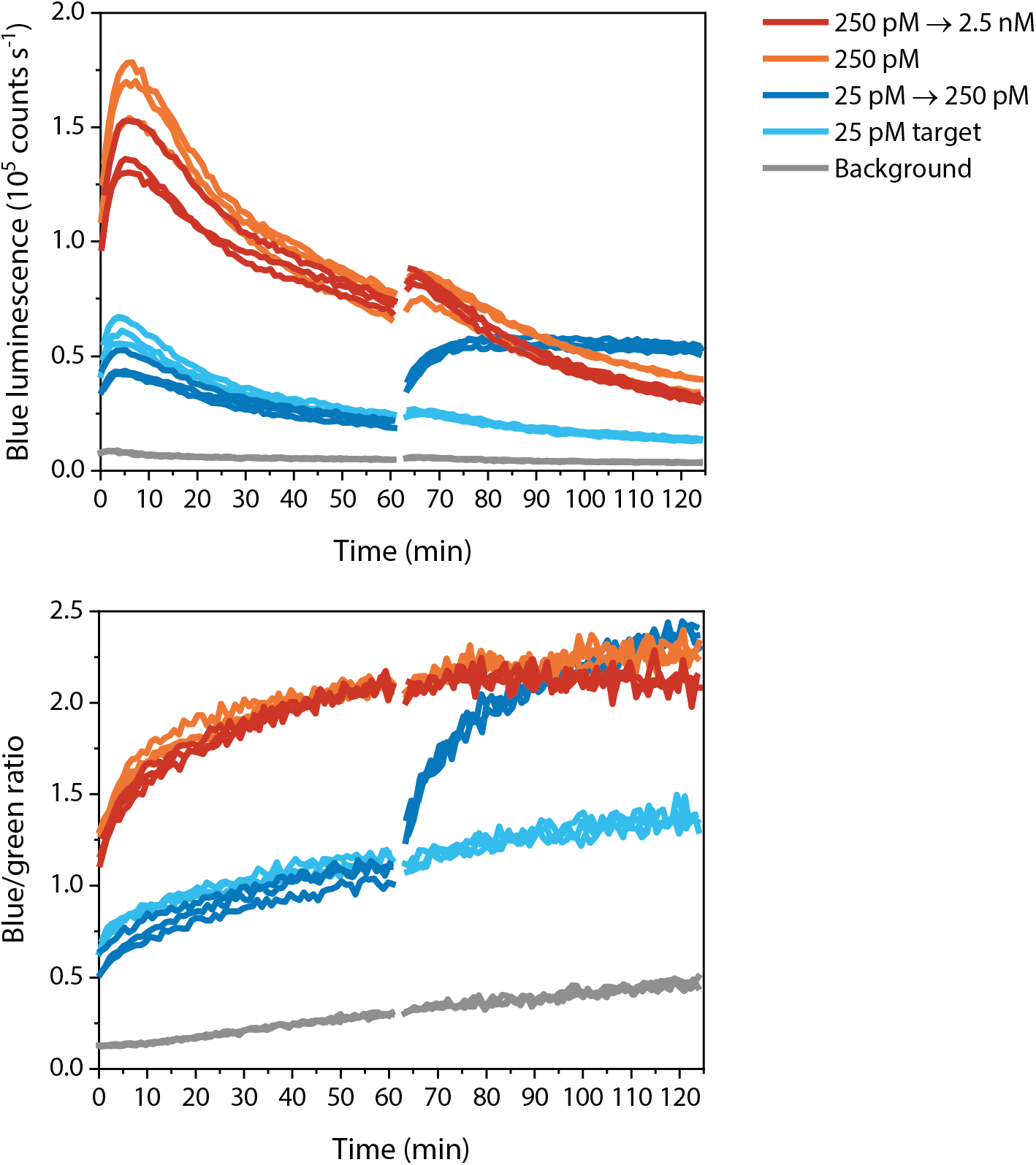
LUNAS kinetics upon increase in target concentration following initial equilibration. dCas9-SB:T7A (1 nM) and dCas9-LB:T7B (1 nM) were incubated with target DNA (30 bp interspace) at indicated concentrations for 1 h, after which the target concentration was increased 10-fold in part of the reactions (red and dark blue lines). The top panel shows the resulting blue luminescence intensity as measured over time. The mNG-NL calibrator luciferase was also included in the reactions (see main text), and the bottom panel shows the blue/green ratio over time. While an increase in target concentration from 25 pM to 250 pM can be seen to result in a rapid increase in blue signal, the increase from 250 pM to 2.5 nM does not result in substantial change in signal compared to that of the control left at 250 pM. This result confirms the extremely slow dissociation rate of dCas9 RNPs, as quick dissociation and redistribution of the dCas9 RNPs would result in a lower signal for 2.5 nM compared to 250 pM target (see Fig. 2C). LUNAS assays were prepared as described for 1-pot RPA-LUNAS assays, excluding primers to preclude RPA, and were performed at 40°C to resemble RPA-LUNAS conditions. Individual replicate traces are shown (n = 3).

