## Supporting Information for "Fast bioluminescent nucleic acid detection using one-pot isothermal amplification and dCas9-based split luciferase complementation"

Supplementary Table 3: Comparison of RT-RPA-LUNAS SARS-CoV-2 assay performance with that of other recent CRISPR diagnostic methods applied to SARS-CoV-2 detection. 22
Supplementary references ..... 23

#### Supplementary Note 1: Thermodynamic model

To shed light on the dependence of the LUNAS response on the various thermodynamic parameters involved, we developed a model of the system (Fig. S1). In this model, the total concentration of the sensor RNP complexes was divided over an active (i.e. DNA-binding competent) and an inactive fraction in a 25:75 ratio, in accordance with previous reports[1,2]. The DNA-binding incompetent sensor complexes (denoted 'Ld' and 'Sd' for the LB and SB complexes respectively) were still included in the model, to account for background signal derived from the split NanoLuc parts of these complexes, but conversion to active complexes (denoted 'La' and 'Sa') and vice versa was excluded. For the interaction of dCas9-SB:gRNA_A and dCas9-LB:gRNA_B with the target DNA (denoted 'T') the same single dissociation equilibrium constant was defined ('$K_DC$'), assuming differences in affinity depending on the exact gRNA/protospacer sequence to be small for functional LUNAS RNP complex pairs. Moreover, dCas9-SB:gRNA_A and dCas9-LB:gRNA_B are considered to bind the target DNA in a non-cooperative fashion, hence dCas9-SB:gRNA_A binds to free target DNA with the same affinity as to target DNA already bound by dCas9-LB:gRNA_B (denoted 'LaT'). Upon formation of the ternary 'LaSaT' complex, the high local concentration (the effective molarity (EM)) of the split NanoLuc fragments promotes complementation, transitioning to the luminescent 'LaSaTa' complex. As non-templated split NanoLuc complementation could also result in luminescence, the total luminescent signal is modelled as the sum of the concentrations of LaSaTa, LaSa, LdSd, LdSa and LaSd, multiplied by a constant.

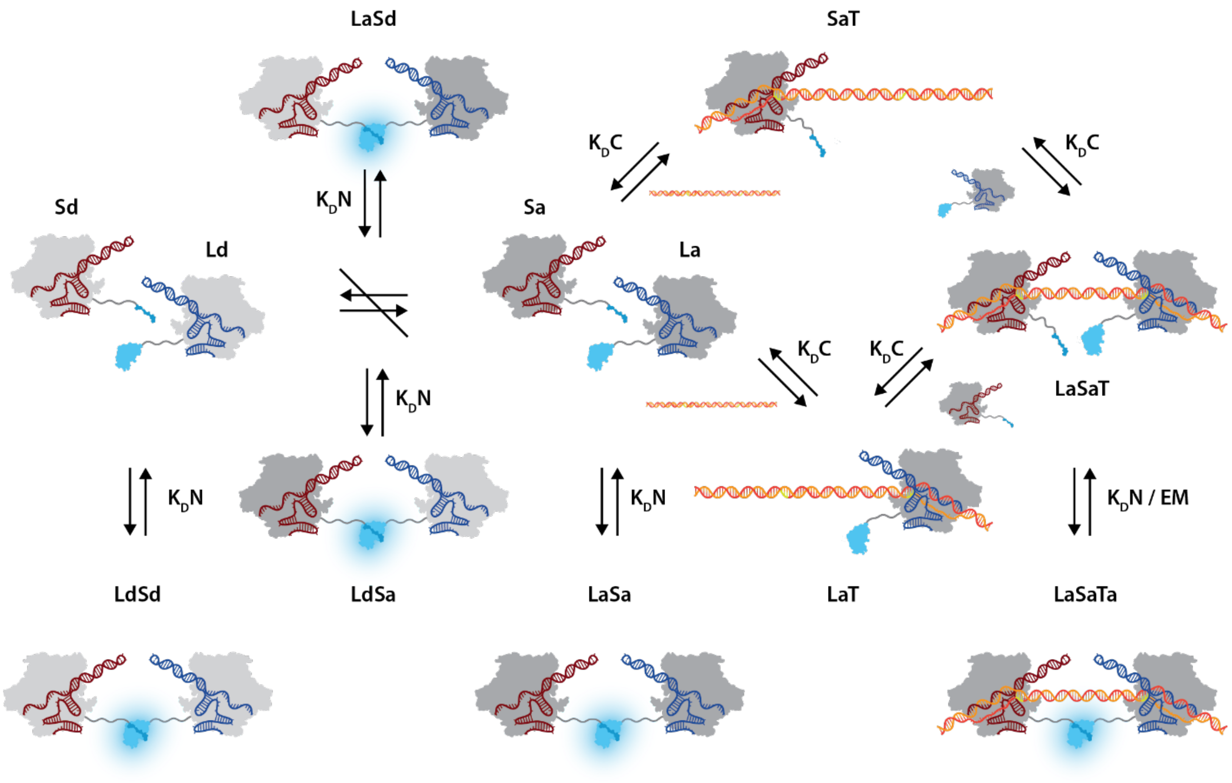

**Supplementary Fig. 1: Model of the thermodynamic interactions in the LUNAS assay.**

The model was implemented using a general framework for equilibrium models developed by Geertjens *et al.*[3], using the following input to build the model in the Python tool (with $K_DN$ = `KDNanoBiT` and $K_DC$ = `KDdCas9`):

```
Equations:

Ld+Sd = LdSd; KDNanoBiT
Ld+Sa = LdSa; KDNanoBiT
La+Sd = LaSd; KDNanoBiT
La+Sa = LaSa; KDNanoBiT
La+T = LaT; KDdCas9
Sa+T = SaT; KDdCas9
LaT+Sa = LaSaT; KDdCas9
SaT+La = LaSaT; KDdCas9
LaSaT = LaSaTa; KDNanoBiT/EM

data_mode: custom
custom_input: constant * (LaSaTa + LaSa + LdSd + LdSa + LaSd)
```

From this, a model was generated in the framework, which is further detailed below for completeness:

The equilibrium concentrations of the dependent species can be determined based on the concentrations of the independent species and the corresponding equilibrium constant, using the following relations:

```
LdSd: Ld*Sd/KDNanoBiT
LdSa: Ld*Sa/KDNanoBiT
LaSd: La*Sd/KDNanoBiT
LaSa: La*Sa/KDNanoBiT
LaT: La*T/KDdCas9
SaT: Sa*T/KDdCas9
LaSaT: La*Sa*T/KDdCas9**2
LaSaTa: EM*La*Sa*T/(KDNanoBiT*KDdCas9**2)
```

The mass balance of the independent species in terms of free and complexed forms:

```
Sa_tot: LaSa + LaSaT + LaSaTa + LdSa + Sa + SaT
La_tot: La + LaSa + LaSaT + LaSaTa + LaSd + LaT
T_tot: LaSaT + LaSaTa + LaT + SaT + T
Sd_tot: LaSd + LdSd + Sd
Ld_tot: Ld + LdSa + LdSd
```

Substituting the relations above in the mass balance equations yields:

```
Sa_tot = EM*La*Sa*T/(KDNanoBiT*KDdCas9**2) + Sa  + Sa*T/KDdCas9 +
La*Sa*T/KDdCas9**2 + La*Sa/KDNanoBiT + Ld*Sa/KDNanoBiT

La_tot = EM*La*Sa*T/(KDNanoBiT*KDdCas9**2) + La  + La*T/KDdCas9 +
La*Sa*T/KDdCas9**2 + La*Sa/KDNanoBiT + La*Sd/KDNanoBiT

T_tot = EM*La*Sa*T/(KDNanoBiT*KDdCas9**2) + T  + La*T/KDdCas9 +
Sa*T/KDdCas9 + La*Sa*T/KDdCas9**2
```

```
Sd_tot = Sd  + La*Sd/KDNanoBiT + Ld*Sd/KDNanoBiT

Ld_tot = Ld  + Ld*Sa/KDNanoBiT + Ld*Sd/KDNanoBiT
```

Finally, these equations were rewritten to equal zero, and divided by the total concentrations on both sides in order to reach a solution faster during solving:

```
Sa: (EM*La*Sa*T/(KDNanoBiT*KDdCas9**2) + Sa - Sa_tot + Sa*T/KDdCas9
+ La*Sa*T/KDdCas9**2 + La*Sa/KDNanoBiT + Ld*Sa/KDNanoBiT) / Sa_tot =
0

La: (EM*La*Sa*T/(KDNanoBiT*KDdCas9**2) + La - La_tot + La*T/KDdCas9
+ La*Sa*T/KDdCas9**2 + La*Sa/KDNanoBiT + La*Sd/KDNanoBiT) / La_tot =
0

T: (EM*La*Sa*T/(KDNanoBiT*KDdCas9**2) + T - T_tot + La*T/KDdCas9 +
Sa*T/KDdCas9 + La*Sa*T/KDdCas9**2) / T_tot = 0

Sd: (Sd - Sd_tot + La*Sd/KDNanoBiT + Ld*Sd/KDNanoBiT) / Sd_tot = 0

Ld: (Ld - Ld_tot + Ld*Sa/KDNanoBiT + Ld*Sd/KDNanoBiT) / Ld_tot = 0
```

This model was fitted to the combined data presented in Fig. 2C using 'KDNanoBiT' = 2.5E-6 M as known parameter[4]. For dCas9:gRNA complex binding to target DNA, an upper limit in $K_D$ of ~0.5 nM was previously reported based on EMSA experiments[5], and here 0.1 nM was taken as an initial value for 'KDdCas9'. The effective molarity of the split NanoLuc fragments (EM) depends on the length and flexibility of the linkers connecting them to the dCas9 proteins, as well as the distance that has to be bridged by the linkers for luciferase complementation. An initial value for the EM was estimated based on the wormlike chain model[6,7]. Considering the 21 residue linkers are made up mostly of GGS repeats, a persistence length of 3.7 Å was assumed[7]. Based on the structure of DNA bound dCas9:gRNA (PDB: 5F9R[8]) and that of NanoLuc (PDB: 5IBO), the distance to be bridged between anchor points (dCas9 C-terminus and SB/LB N-terminus) for the 50 bp interspace target used is roughly 49 Å. From this, we estimated an EM of ~ 31 µM[6]. For the factor converting luminescent entity concentrations to luminescence signal intensity, the initial value was set as 'constant' = 1E+15. The following parameter estimates were obtained (fitted lines shown in Fig. 2C):

```
EM = 1.048e-05 M
KDdCas9 = 1.767e-11 M
constant = 1.939e+15

Root Mean Squared Error (10 nM dCas9-SB RNP + 1 nM dCas9-LB RNP) =
2.01e+04
Root Mean Squared Error (1 nM dCas9-SB RNP + 1 nM dCas9-LB RNP) =
2.02e+04
```

$R^2$ (10 nM dCas9-SB RNP + 1 nM dCas9-LB RNP)= 0,982
$R^2$ (1 nM dCas9-SB RNP + 1 nM dCas9-LB RNP)= 0,955

To gauge the dependence of the signal response on tunable sensor parameters, we performed simulations in which the concentration ratio of sensor RNP complexes was varied, and we simulated the effect of different split NanoLuc binding affinities (Fig. S2). For this, parameter estimates described above were used.

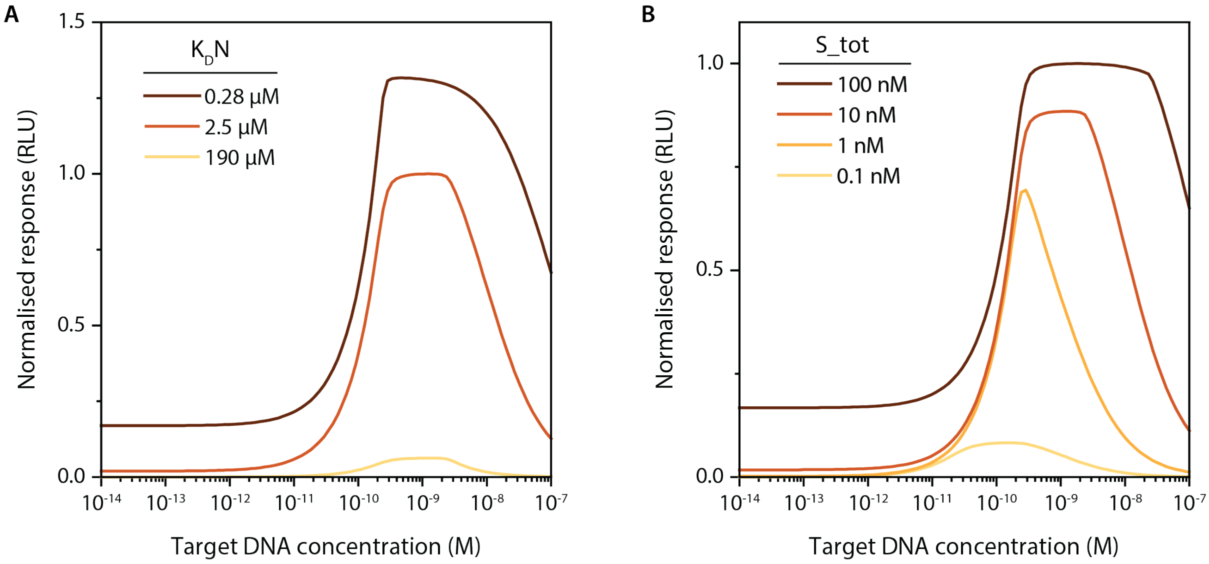

**Supplementary Fig. 2: Model simulations of LUNAS response for a variation in sensor parameters. A** Varying the affinity of small BiT (SB) for binding to large BiT (LB) ($K_DN$). **B** Varying the total concentration of dCas9-SB:gRNA_A (S_tot), keeping total dCas9-LB:gRNA_B at 1 nM. For both **(A)** and **(B)**, the response was normalised to the maximum of the orange line, corresponding to $K_DN$ = 2.5 µM and S_tot = 10 nM. Other parameters were set as follows: EM = 10.48 µM, $K_dC$ = 17.67 pM, constant = $1.939 \times 10^{15}$. For **(B)**, $K_DN$ = 2.5 µM. For **(A)**, L_tot = 1 nM and S_tot = 10 nM.

### Supplementary Note 2: Protein coding sequences and translations

**dCas9-SB / dCas9-LB**

The following combined sequence codes for dCas9-SB (default) and dCas9-LB (after SpeI restriction digest and subsequent self-ligation of the large fragment). Colour codes: dCas9 highlighted in yellow; SB in cyan; LB in green; SpeI restriction site in magenta, Strep-tag II in red font.

```
atggataagaaatactcaataggcttagctatcggcacaaatagcgtcggatgggcggtg
 M   D   K   K   Y   S   I   G   L   A   I   G   T   N   S   V   G   W   A   V
atcactgatgaatataaggttccgtctaaaaagttcaaggttctgggaaatacagaccgc
 I   T   D   E   Y   K   V   P   S   K   K   F   K   V   L   G   N   T   D   R
cacagtatcaaaaaaaatcttataggggctcttttatttgacagtggagagacagcggaa
 H   S   I   K   K   N   L   I   G   A   L   L   F   D   S   G   E   T   A   E
gcgactcgtctcaaacggacagctcgtagaaggtatacacgtcggaagaatcgtatttgt
 A   T   R   L   K   R   T   A   R   R   R   Y   T   R   R   K   N   R   I   C
tatctacaggagattttttcaaatgagatggcgaaagtagatgatagtttctttcatcga
 Y   L   Q   E   I   F   S   N   E   M   A   K   V   D   D   S   F   F   H   R
cttgaagagtcttttttggtggaagaagacaagaagcatgaacgtcatcctatttttgga
 L   E   E   S   F   L   V   E   E   D   K   K   H   E   R   H   P   I   F   G
aatatagtagatgaagttgcttatcatgagaaatatccaactatctatcatctgcgaaaa
 N   I   V   D   E   V   A   Y   H   E   K   Y   P   T   I   Y   H   L   R   K
aaattggtagattctactgataaagcggatttgcgcttaatctatttggccttagcgcat
 K   L   V   D   S   T   D   K   A   D   L   R   L   I   Y   L   A   L   A   H
atgattaagtttcgtggtcattttttgattgagggagatttaaatcctgataatagtgat
 M   I   K   F   R   G   H   F   L   I   E   G   D   L   N   P   D   N   S   D
gtggacaaactatttatccagttggtacaaacctacaatcaattatttgaagaaaaccct
 V   D   K   L   F   I   Q   L   V   Q   T   Y   N   Q   L   F   E   E   N   P
attaacgcaagtggagtagatgctaaagcgattctttctgcacgattgagtaaatcaaga
 I   N   A   S   G   V   D   A   K   A   I   L   S   A   R   L   S   K   S   R
cgattagaaaatctcattgctcagctccccggtgagaagaaaaatggcttatttgggaat
 R   L   E   N   L   I   A   Q   L   P   G   E   K   K   N   G   L   F   G   N
ctcattgctttgtcattgggtttgacccctaattttaaatcaaattttgatttggcagaa
 L   I   A   L   S   L   G   L   T   P   N   F   K   S   N   F   D   L   A   E
gatgctaaattacagctttcaaaagatacttacgatgatgatttagataatttattggcg
 D   A   K   L   Q   L   S   K   D   T   Y   D   D   D   L   D   N   L   L   A
caaattggagatcaatatgctgatttgtttttggcagctaagaatttatcagatgctatt
 Q   I   G   D   Q   Y   A   D   L   F   L   A   A   K   N   L   S   D   A   I
ttactttcagatatcctaagagtaaatactgaaataactaaggctcccctatcagcttca
 L   L   S   D   I   L   R   V   N   T   E   I   T   K   A   P   L   S   A   S
atgattaaacgctacgatgaacatcatcaagacttgactcttttaaaagctttagttcga
 M   I   K   R   Y   D   E   H   H   Q   D   L   T   L   L   K   A   L   V   R
caacaacttccagaaaagtataaagaaatcttttttgatcaatcaaaaaacggatatgca
 Q   Q   L   P   E   K   Y   K   E   I   F   F   D   Q   S   K   N   G   Y   A
ggttatattgatgggggagctagccaagaagaattttataaatttatcaaaccaatttta
 G   Y   I   D   G   G   A   S   Q   E   E   F   Y   K   F   I   K   P   I   L
gaaaaaatggatggtactgaggaattattggtgaaactaaatcgtgaagatttgctgcgc
 E   K   M   D   G   T   E   E   L   L   V   K   L   N   R   E   D   L   L   R
aagcaacggacctttgacaacggctctattccccatcaaattcacttgggtgagctgcat
 K   Q   R   T   F   D   N   G   S   I   P   H   Q   I   H   L   G   E   L   H
gctattttgagaagacaagaagacttttatccatttttaaaagacaatcgtgagaagatt
 A   I   L   R   R   Q   E   D   F   Y   P   F   L   K   D   N   R   E   K   I
gaaaaaatcttgacttttcgaattccttattatgttggtccattggcgcgtggcaatagt
 E   K   I   L   T   F   R   I   P   Y   Y   V   G   P   L   A   R   G   N   S
cgttttgcatggatgactcggaagtctgaagaaacaattaccccatggaattttgaagaa
 R   F   A   W   M   T   R   K   S   E   E   T   I   T   P   W   N   F   E   E
```

gttgtcgataaaggtgcttcagctcaatcatttattgaacgcatgacaaactttgataaa

V V D K G A S A Q S F I E R M T N F D K

aatcttccaaatgaaaaagtactaccaaaacatagtttgctttatgagtattttacggtt

N L P N E K V L P K H S L L Y E Y F T V

tataacgaattgacaaaggtcaaatatgttactgaaggaatgcgaaaaccagcatttctt

Y N E L T K V K Y V T E G M R K P A F L

tcaggtgaacagaagaaagccattgttgatttactcttcaaaacaaatcgaaaagtaacc

S G E Q K K A I V D L L F K T N R K V T

gttaagcaattaaaagaagattatttcaaaaaaatagaatgttttgatagtgttgaaatt

V K Q L K E D Y F K K I E C F D S V E I

tcaggagttgaagatagatttaatgcttcattaggtacctaccatgatttgctaaaaatt

S G V E D R F N A S L G T Y H D L L K I

attaaagataaagatttttggataatgaagaaaatgaagatatcttagaggatattgtt

I K D K D F L D N E E N E D I L E D I V

ttaacattgaccttatttgaagatagggagatgattgaggaaagacttaaaacatatgct

L T L T L F E D R E M I E E R L K T Y A

cacctctttgatgataaggtgatgaaacagcttaaacgtcgccgttatactggttggga

H L F D D K V M K Q L K R R R Y T G W G

cgtttgtctcgaaaattgattaatggtattagggataagcaatctggcaaaacaatatta

R L S R K L I N G I R D K Q S G K T I L

gatttttgaaatcagatggttttgccaatcgcaattttatgcagctgatccatgatgat

D F L K S D G F A N R N F M Q L I H D D

agtttgacatttaaagaagacattcaaaaagcacaagtgtctggacaaggcgatagttta

S L T F K E D I Q K A Q V S G Q G D S L

catgaacatattgcaaatttagctggtagccctgctattaaaaaaggtattttacagact

H E H I A N L A G S P A I K K G I L Q T

gtaaaagttgttgatgaattggtcaaagtaatggggcggcataagccagaaaatatcgtt

V K V V D E L V K V M G R H K P E N I V

attgaaatggcacgtgaaaatcagacaactcaaaagggccagaaaaattcgcgagagcgt

I E M A R E N Q T T Q K G Q K N S R E R

atgaaacgaatcgaagaaggtatcaaagaattaggaagtcagattcttaaagagcatcct

M K R I E E G I K E L G S Q I L K E H P

gttgaaaatactcaattgcaaaatgaaaagctctatctctattatctccaaaatggaaga

V E N T Q L Q N E K L Y L Y Y L Q N G R

gacatgtatgtggaccaagaattagatattaatcgtttaagtgattatgatgtcgatgcc

D M Y V D Q E L D I N R L S D Y D V D A

attgttccacaaagtttccttaaagacgattcaatagacaataaggtcttaacgcgttct

I V P Q S F L K D D S I D N K V L T R S

gataaaaatcgtggtaaatcggataacgttccaagtgaagaagtagtcaaaaagatgaaa

D K N R G K S D N V P S E E V V K K M K

aactattggagacaacttctaaacgccaagttaatcactcaacgtaagtttgataattta

N Y W R Q L L N A K L I T Q R K F D N L

acgaaagctgaacgtggaggtttgagtgaacttgataaagctggttttatcaaacgccaa

T K A E R G G L S E L D K A G F I K R Q

ttggttgaaactcgccaaatcactaagcatgtggcacaaattttggatagtcgcatgaat

L V E T R Q I T K H V A Q I L D S R M N

actaaatacgatgaaaatgataaacttattcgagaggttaaagtgattaccttaaaatct

T K Y D E N D K L I R E V K V I T L K S

aaattagtttctgacttccgaaaagatttccaattctataaagtacgtgagattaacaat

K L V S D F R K D F Q F Y K V R E I N N

taccatcatgcccatgatgcgtatctaaatgccgtcgttggaactgctttgattaagaaa

Y H H A H D A Y L N A V V G T A L I K K

tatccaaaacttgaatcggagtttgtctatggtgattataaagtttatgatgttcgtaaa

Y P K L E S E F V Y G D Y K V Y D V R K

atgattgctaagtctgagcaagaaataggcaaagcaaccgcaaaatatttcttttactct

M I A K S E Q E I G K A T A K Y F F Y S

aatatcatgaacttcttcaaaacagaaattacacttgcaaatggagagattcgcaaacgc

N I M N F F K T E I T L A N G E I R K R

```
cctctaatcgaaactaatggggaaactggagaaattgtctgggataaagggcgagatttt
 P  L  I  E  T  N  G  E  T  G  E  I  V  W  D  K  G  R  D  F
gccacagtgcgcaaagtattgtccatgccccaagtcaatattgtcaagaaaacagaagta
 A  T  V  R  K  V  L  S  M  P  Q  V  N  I  V  K  K  T  E  V
cagacaggcggattctccaaggagtcaattttaccaaaaagaaattcggacaagcttatt
 Q  T  G  G  F  S  K  E  S  I  L  P  K  R  N  S  D  K  L  I
gctcgtaaaaaagactgggatccaaaaaaatatggtggttttgatagtccaacggtagct
 A  R  K  K  D  W  D  P  K  K  Y  G  G  F  D  S  P  T  V  A
tattcagtcctagtggttgctaaggtggaaaaagggaaatcgaagaagttaaaatccgtt
 Y  S  V  L  V  V  A  K  V  E  K  G  K  S  K  K  L  K  S  V
aaagagttactagggatcacaattatggaaagaagttcctttgaaaaaaatccgattgac
 K  E  L  L  G  I  T  I  M  E  R  S  S  F  E  K  N  P  I  D
tttttagaagctaaaggatataaggaagttaaaaaagacttaatcattaaactacctaaa
 F  L  E  A  K  G  Y  K  E  V  K  K  D  L  I  I  K  L  P  K
tatagtctttttgagttagaaaacggtcgtaaacggatgctggctagtgccggagaatta
 Y  S  L  F  E  L  E  N  G  R  K  R  M  L  A  S  A  G  E  L
caaaaaggaaatgagctggctctgccaagcaaatatgtgaatttttatatttagctagt
 Q  K  G  N  E  L  A  L  P  S  K  Y  V  N  F  L  Y  L  A  S
cattatgaaaagttgaagggtagtccagaagataacgaacaaaaacaattgtttgtggag
 H  Y  E  K  L  K  G  S  P  E  D  N  E  Q  K  Q  L  F  V  E
cagcataagcattatttagatgagattattgagcaaatcagtgaattttctaagcgtgtt
 Q  H  K  H  Y  L  D  E  I  I  E  Q  I  S  E  F  S  K  R  V
attttagcagatgccaatttagataaagttcttagtgcatataacaaacatagagacaaa
 I  L  A  D  A  N  L  D  K  V  L  S  A  Y  N  K  H  R  D  K
ccaatacgtgaacaagcagaaaatattattcatttatttacgttgacgaatcttggagct
 P  I  R  E  Q  A  E  N  I  I  H  L  F  T  L  T  N  L  G  A
cccgctgcttttaaatattttgatacaacaattgatcgtaaacgatatacgtctacaaaa
 P  A  A  F  K  Y  F  D  T  T  I  D  R  K  R  Y  T  S  T  K
gaagttttagatgccactcttatccatcaatccatcactggtctttatgaaacacgcatt
 E  V  L  D  A  T  L  I  H  Q  S  I  T  G  L  Y  E  T  R  I
gatttgagtcagctaggaggtgacaccggtgggggtagcggcggctcgggggggtagtggt
 D  L  S  Q  L  G  G  D  T  G  G  G  S  G  G  S  G  G  S  G
ggaagcgggggttcaaagcttactagtgttaccggctatcgtctgtttgaaaaagagagc
 G  S  G  G  S  K  L  T  S  V  T  G  Y  R  L  F  E  K  E  S
ggatccggtggaagctggagccatccgcagtttgaaaaataaactagtgtcttcacactc
 G  S  G  G  S  W  S  H  P  Q  F  E  K  -  T  S  V  F  T  L
gaagatttcgttggggactgggaacagacagccgcctacaacctggaccaagtccttgaa
 E  D  F  V  G  D  W  E  Q  T  A  A  Y  N  L  D  Q  V  L  E
cagggaggtgtgtccagtttgctgcagaatctcgccgtgtccgtaactccgatccaaagg
 Q  G  G  V  S  S  L  L  Q  N  L  A  V  S  V  T  P  I  Q  R
attgtccggagcggtgaaaatgccctgaagatcgacatccatgtcatcatcccgtatgaa
 I  V  R  S  G  E  N  A  L  K  I  D  I  H  V  I  I  P  Y  E
ggtctgagcgccgaccaaatggcccagatcgaagaggtgtttaaggtggtgtaccctgtg
 G  L  S  A  D  Q  M  A  Q  I  E  E  V  F  K  V  V  Y  P  V
gatgatcatcactttaaggtgatcctgccctatggcacactggtaatcgacggggttacg
 D  D  H  H  F  K  V  I  L  P  Y  G  T  L  V  I  D  G  V  T
ccgaacatgctgaactatttcggacggccgtatgaaggcatcgccgtgttcgacggcaaa
 P  N  M  L  N  Y  F  G  R  P  Y  E  G  I  A  V  F  D  G  K
aagatcactgtaacagggaccctgtggaacggcaacaaaattatcgacgagcgcctgatc
 K  I  T  V  T  G  T  L  W  N  G  N  K  I  I  D  E  R  L  I
acccccgacggctccatgctgttccgagtaaccatcaacagcggtggaagctggagccat
 T  P  D  G  S  M  L  F  R  V  T  I  N  S  G  G  S  W  S  H
ccgcagtttgaaaaataa
 P  Q  F  E  K  -
```

**mNG-NL calibrator luciferase**

Coding and amino acid sequences for mNeongreen-NanoLuc fusion protein[9,10]. Colour codes: mNeonGreen-ΔC10 in green; NanoLuc-ΔN5 in cyan; His-tag in blue font; StrepTag in red font.

```
atgggcagcagccatcatcatcatcatcacagcagcggcctggtgccgcgcggcagccat
 M   G   S   S   H   H   H   H   H   H   S   S   G   L   V   P   R   G   S   H
atggtaagtaaaggtgaagaagacaatatggcttctctgcctgccacacatgagcttcat
 M   V   S   K   G   E   E   D   N   M   A   S   L   P   A   T   H   E   L   H
atttttgggagcataaacggagtggatttcgacatggtaggtcagggtacggggaaccct
 I   F   G   S   I   N   G   V   D   F   D   M   V   G   Q   G   T   G   N   P
aacgatggatatgaggagttgaatcttaaaagcacaaagggtgatctgcagttctcgccc
 N   D   G   Y   E   E   L   N   L   K   S   T   K   G   D   L   Q   F   S   P
tggatcctggtgccgcatataggttatggtttccatcagtatcttccatacccggatggc
 W   I   L   V   P   H   I   G   Y   G   F   H   Q   Y   L   P   Y   P   D   G
atgagccctttttcaggccgcaatggtagatggctcaggatatcaagtgcatcggaccatg
 M   S   P   F   Q   A   A   M   V   D   G   S   G   Y   Q   V   H   R   T   M
cagtttgaagatggggcgtctttgacggtaaattacaggtacacctatgagggtagccat
 Q   F   E   D   G   A   S   L   T   V   N   Y   R   Y   T   Y   E   G   S   H
ataaagggagaagcgcaggtgaagggaactggattcccagcggatggcccagtcatgaca
 I   K   G   E   A   Q   V   K   G   T   G   F   P   A   D   G   P   V   M   T
aacagcctcaccgctgctgattggtgccgatccaagaaaacgtatccaaacgataaaact
 N   S   L   T   A   A   D   W   C   R   S   K   K   T   Y   P   N   D   K   T
atcatttctacttttaagtggtcctatacaacaggaaacgggaaacgctatcgttcaacg
 I   I   S   T   F   K   W   S   Y   T   T   G   N   G   K   R   Y   R   S   T
gcccgcacgacctacacgtttgcaaagccaatggctgcgaattatctgaaaaaccagccg
 A   R   T   T   Y   T   F   A   K   P   M   A   A   N   Y   L   K   N   Q   P
atgtatgtgttccgtaaaaccgaactgaaacattctaaaacggagctcaatttcaaggaa
 M   Y   V   F   R   K   T   E   L   K   H   S   K   T   E   L   N   F   K   E
tggcagaaggcatttaccggttttgaagatttcgtgggtgattggcgacaaacggccggt
 W   Q   K   A   F   T   G   F   E   D   F   V   G   D   W   R   Q   T   A   G
tacaatttggatcaggtgttagaacaaggggggcgtaagctccctgttccagaatttagga
 Y   N   L   D   Q   V   L   E   Q   G   G   V   S   S   L   F   Q   N   L   G
gtgagcgtgacacctattcagcgcattgtgctgagcggcgaaaatggtctgaaaattgat
 V   S   V   T   P   I   Q   R   I   V   L   S   G   E   N   G   L   K   I   D
attcatgtgatcatcccttacgaaggcctgtctggggatcaaatgggacagattgaaaaa
 I   H   V   I   I   P   Y   E   G   L   S   G   D   Q   M   G   Q   I   E   K
atcttcaaagtagtttatccggtcgacgatcatcattttaaagtaattctgcactatggg
 I   F   K   V   V   Y   P   V   D   D   H   H   F   K   V   I   L   H   Y   G
acactcgttatcgatggagtcacgccgaatatgatagactacttcggtcgcccgtacgaa
 T   L   V   I   D   G   V   T   P   N   M   I   D   Y   F   G   R   P   Y   E
ggaatcgcggttttcgatggaaaaaaaatcacagtaacgggcacattgtggaacgggaat
 G   I   A   V   F   D   G   K   K   I   T   V   T   G   T   L   W   N   G   N
aaaatcatagacgaacgcctcattaaccctgatggatctttactgttccgcgtcacaatt
 K   I   I   D   E   R   L   I   N   P   D   G   S   L   L   F   R   V   T   I
aatggcgttacaggttggcgactgtgtgaacgtattctcgcaggtaccacatctgcgtgg
 N   G   V   T   G   W   R   L   C   E   R   I   L   A   G   T   T   S   A   W
agccatcctcagttcgaaaaataa
 S   H   P   Q   F   E   K   -
```

#### Supplementary Figures

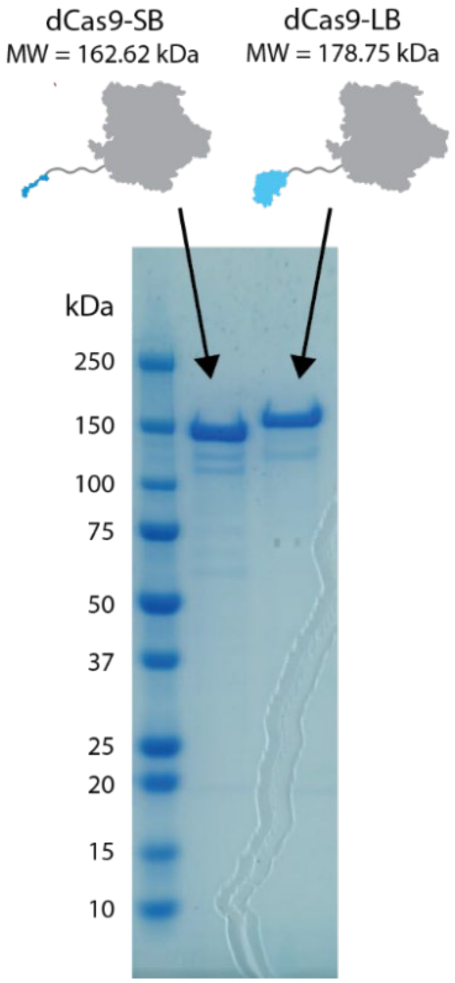

**Supplementary Fig. 3: Expression and purification of LUNAS proteins.** Reducing SDS-PAGE (4-20%) analysis of dCas9-SB and dCas9-LB. After expression in *E. coli* BL21, the proteins were purified by Strep-Tactin XT chromatography.

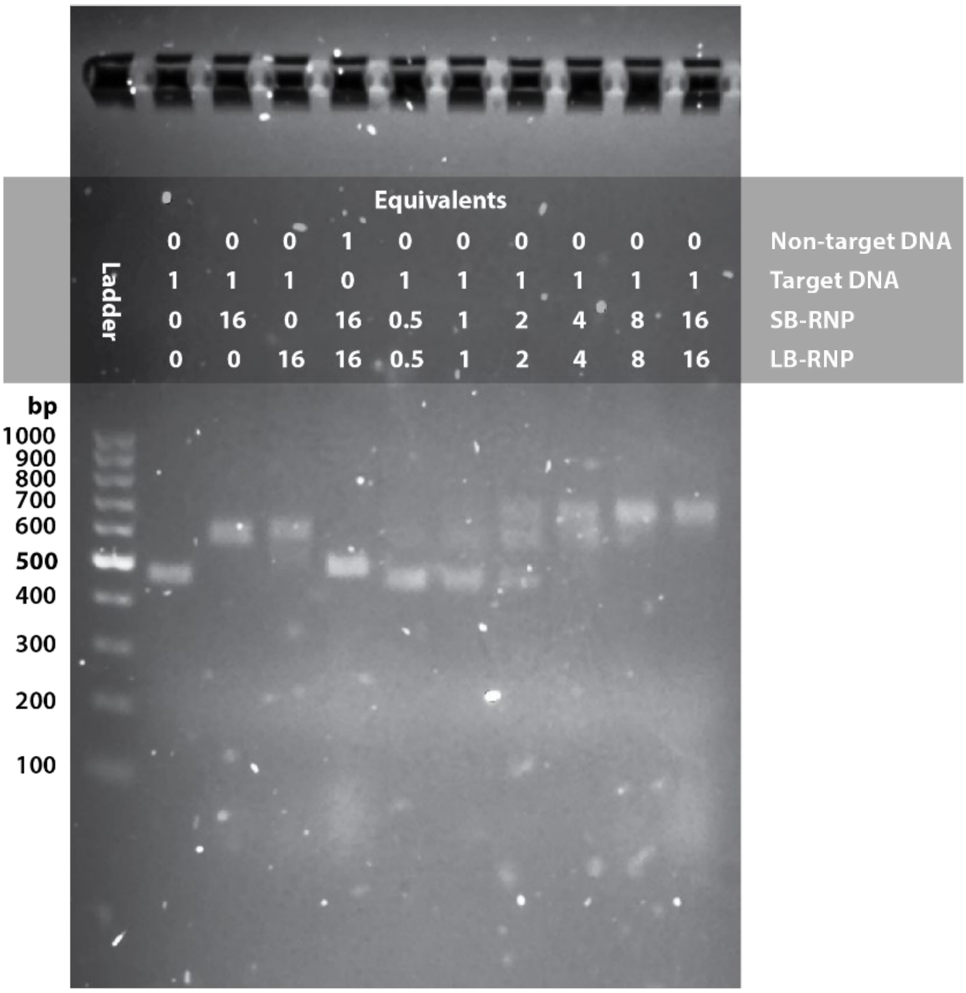

**Supplementary Fig. 4: Electrophoretic mobility shift assay (EMSA) with LUNAS RNPs.** 12.5 nM of non-target (510 bp) or target (473 bp, 30 bp interspace) dsDNA fragment was incubated with 0.5 to 16 equivalents of RNP-SB and/or RNP-LB for 1 hour at RT in LUNAS RNP buffer. Reactions were loaded on a 2% agarose gel containing 1x SYBR safe and run for 55 minutes at 100V in 1x TAE buffer.

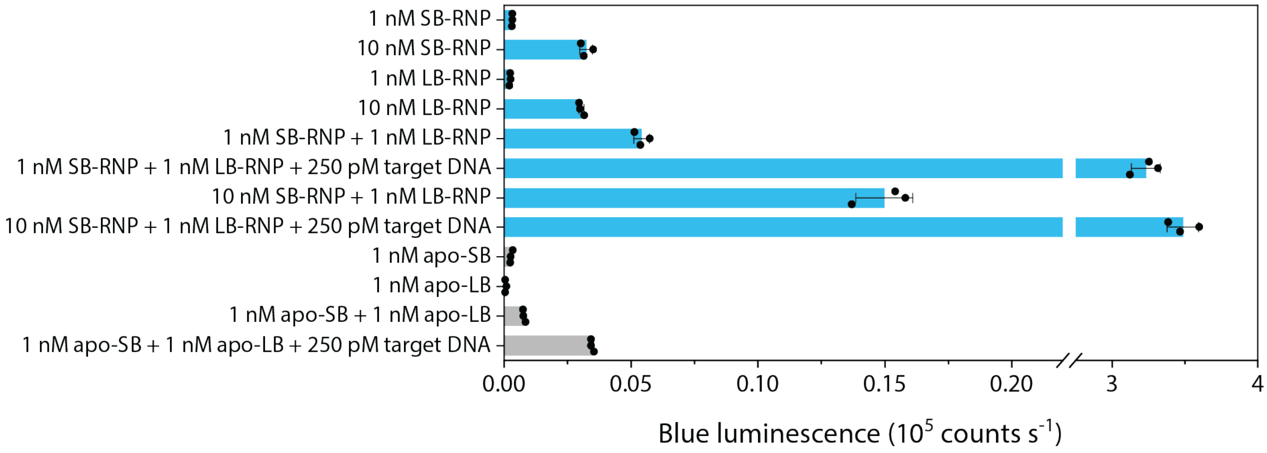

**Supplementary Fig. 5: LUNAS background luminescence**. Individual RNPs show low levels of luminescence, and the background signal observed in LUNAS assays appears to be mostly resulting from split-NanoLuc complementation upon target-independent binding of SB-RNP and LB-RNP to each other. Using 10 nM SB-RNP instead of 1 nM SB-RNP results in roughly 3-fold higher background signal. Apo proteins (i.e. without gRNA) also show low luminescence, which does not increase above LUNAS background levels upon presence of DNA, confirming that gRNA-guided specific target binding is required for strong increase in luminescence. Bars represent means, error bars show SD and individual replicates ($n$ = 3) are represented as dots.

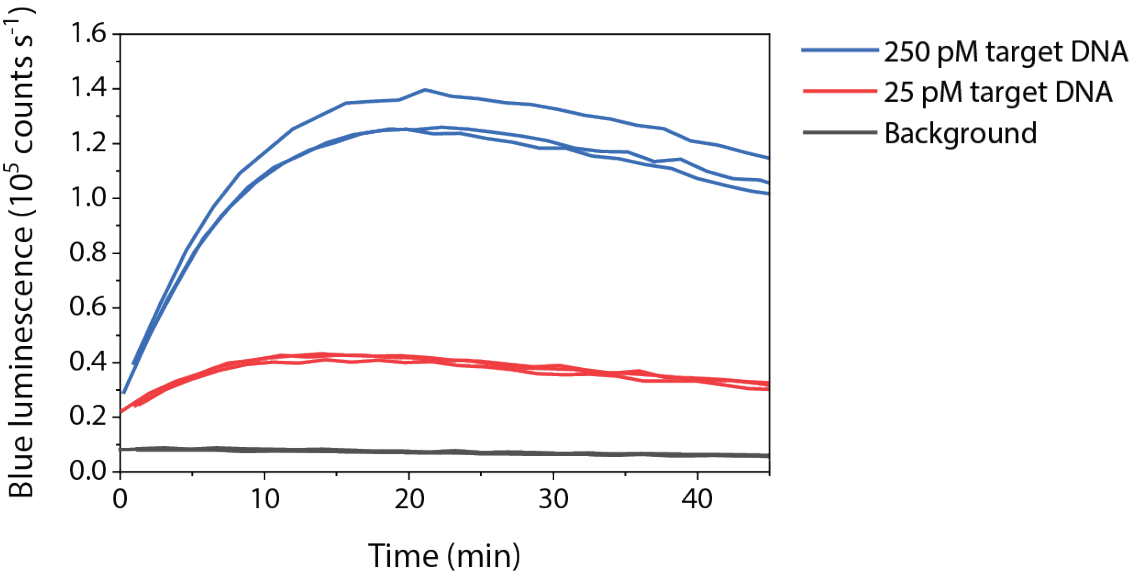

**Supplementary Fig. 6: Kinetics of intensiometric LUNAS.** 1 nM dCas9-SB:gRNA_T7A and 1 nM dCas9-LB:gRNA_T7B were combined with target (30 bp interspace) and NanoGlo substrate (1000-fold final dilution) directly before start of measurement over time. Individual replicate traces (n = 3) are shown.

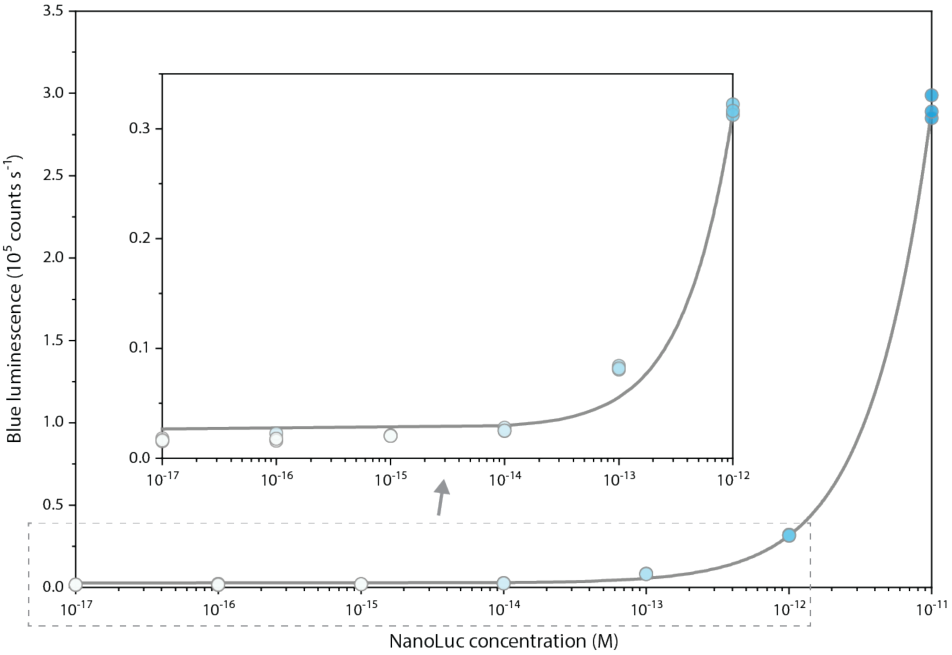

**Supplementary Fig. 7: NanoLuc titration curve.** A 10-fold serial dilution series of NanoLuc in LUNAS RNP buffer was combined with NanoGlo substrate (2000-fold final dilution) and blue luminescence intensity was measured. The inset zooms in on the portion of the main graph indicated in the dashed box. This data shows that the minimal active NanoLuc concentration that can be detected under LUNAS conditions in 20 µL is in the 10 – 100 fM range. Since complemented split-NanoLuc (NanoBiT) has a relative luciferase activity of ~ 37% compared to that of full-length NanoLuc, the minimal concentration of complemented split-NanoLuc that can be detected under these conditions is presumably on the order of ~100 fM[4]. Individual replicates (*n* = 3) are shown as circles, the line represents a linear fit to the full data range (Pearson's r = 0.99969; $R^2$ (COD) = 0.99938).

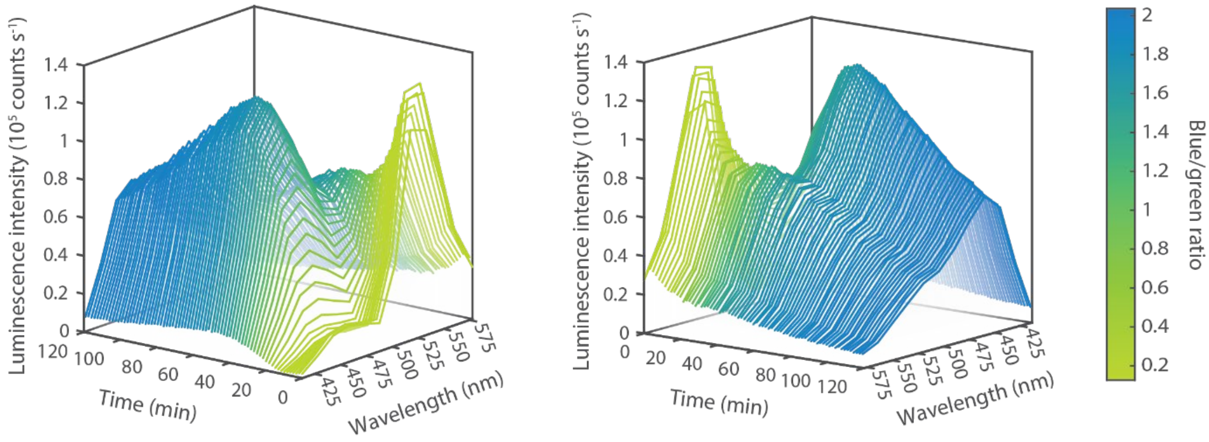

**Supplementary Fig. 8: Ratiometric RPA-LUNAS luminescence spectra over time.** Luminescence spectra of a single SARS-CoV-2 RT-RPA-LUNAS reaction replicate (from Figure 4D, 200 cp input) as measured over time (extended data version of Figure 4C). The right waterfall graph shows the backside of the left graph. The rapid increase in blue signal can be observed, followed by an overall gradually decreasing luminescence due to substrate depletion, which can be observed for the green signal already from the start. However, the blue/green ratio stays relatively constant over time after the initial rise in blue LUNAS signal.

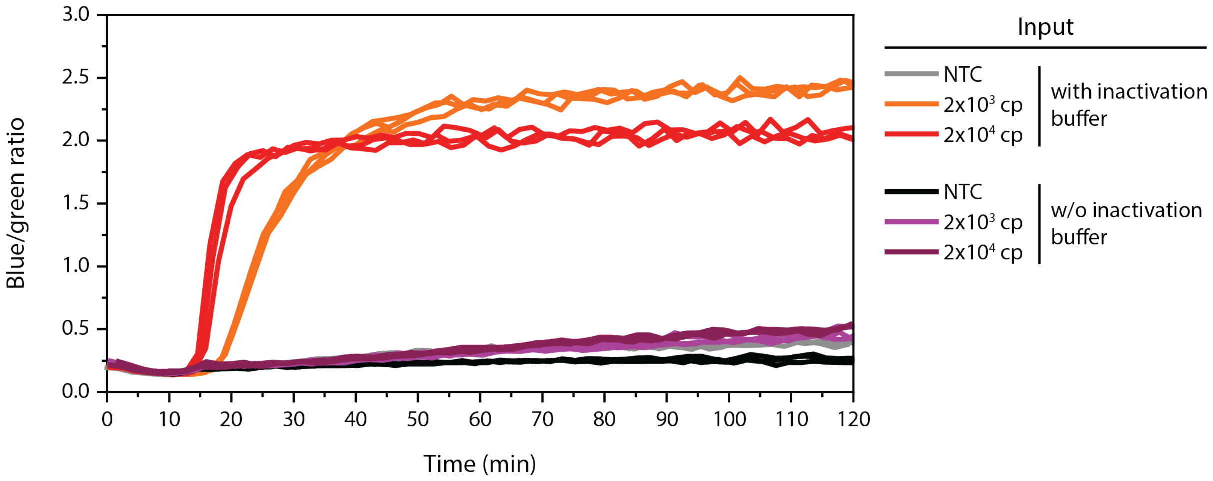

**Supplementary Fig. 9: Importance of RNase inactivation for RT-RPA-LUNAS in saliva samples.** The 1-pot RT-RPA-LUNAS SARS-CoV-2 assay was performed for mock Covid-19 saliva samples pretreated with or without the RNase inactivation buffer. Both pure saliva and saliva 1:1 diluted in inactivation buffer (100 mM TCEP, 1mM EDTA, 1U/µL murine RNase inhibitor, 10 mM Tris-HCl, pH 8.0) were spiked with IVT SARS-CoV-2 target RNA fragment and then heated for 5 minutes at 95°C. After a brief cool-down, these samples were added to ratiometric RT-RPA-LUNAS reactions. Clearly, only the samples treated with the inactivation buffer show a clear LUNAS response. Individual replicate traces are shown (*n* = 3).

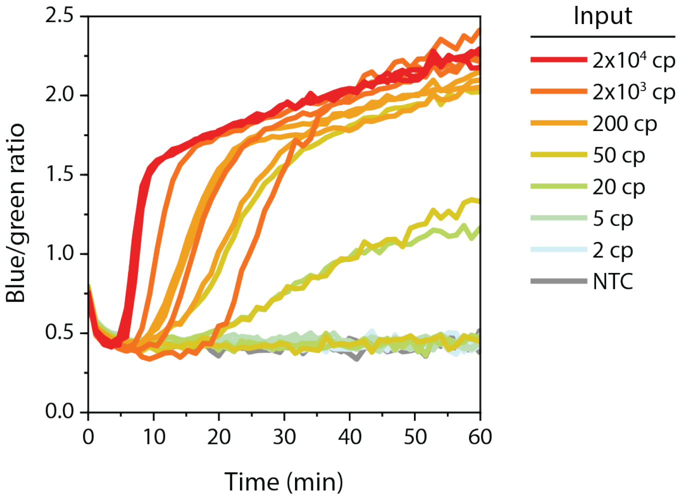

**Supplementary Fig. 10: Camera-based RT-RPA-LUNAS for SARS-CoV-2 RNA detection from saliva.** Camera-based readout of experiment that is similar to the one shown in Fig. 4F with saliva input, with ratiometric RT-RPA-LUNAS response extracted from pictures recorded by camera. Individual replicate traces ($n$ = 3) are shown.

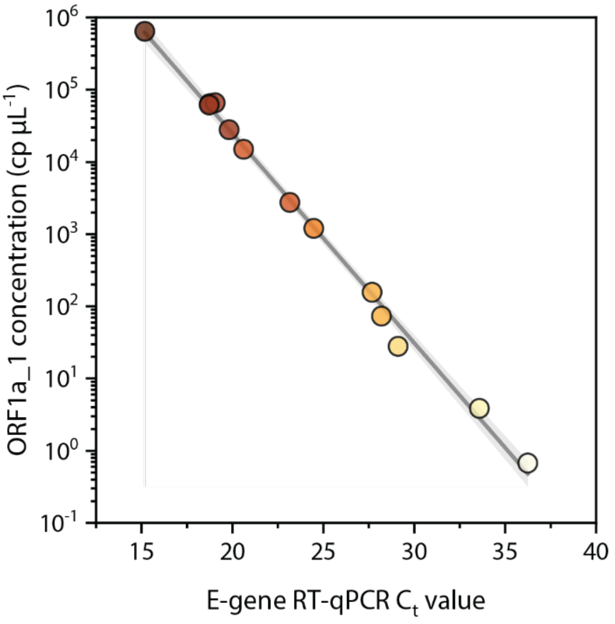

| | |
|---|---|
| Equation | $\log_{10}(y) = a + b*x$ |
| a | 10.19528 ± 0.14663 |
| b | -0.29009 ± 0.00587 |
| Residual Sum of Squares | 0.18477 |
| Pearson's r | -0.99775 |
| R-Square (COD) | 0.99551 |
| Adj. R-Square | 0.9951 |

**Supplementary Fig. 11: RT-ddPCR quantification of SARS-CoV-2 RNA in clinical samples.** ORF1a_1 concentration quantitated by ddPCR versus corresponding E-gene RT-qPCR $C_t$ value. A linear regression model was fitted to the log-transformed data (grey line, with 95% confidence bands), parameters are shown in the table on the right. Single replicates were measured and are represented as filled circles. The reported concentrations correspond to the RNA concentrations in the extraction eluates.

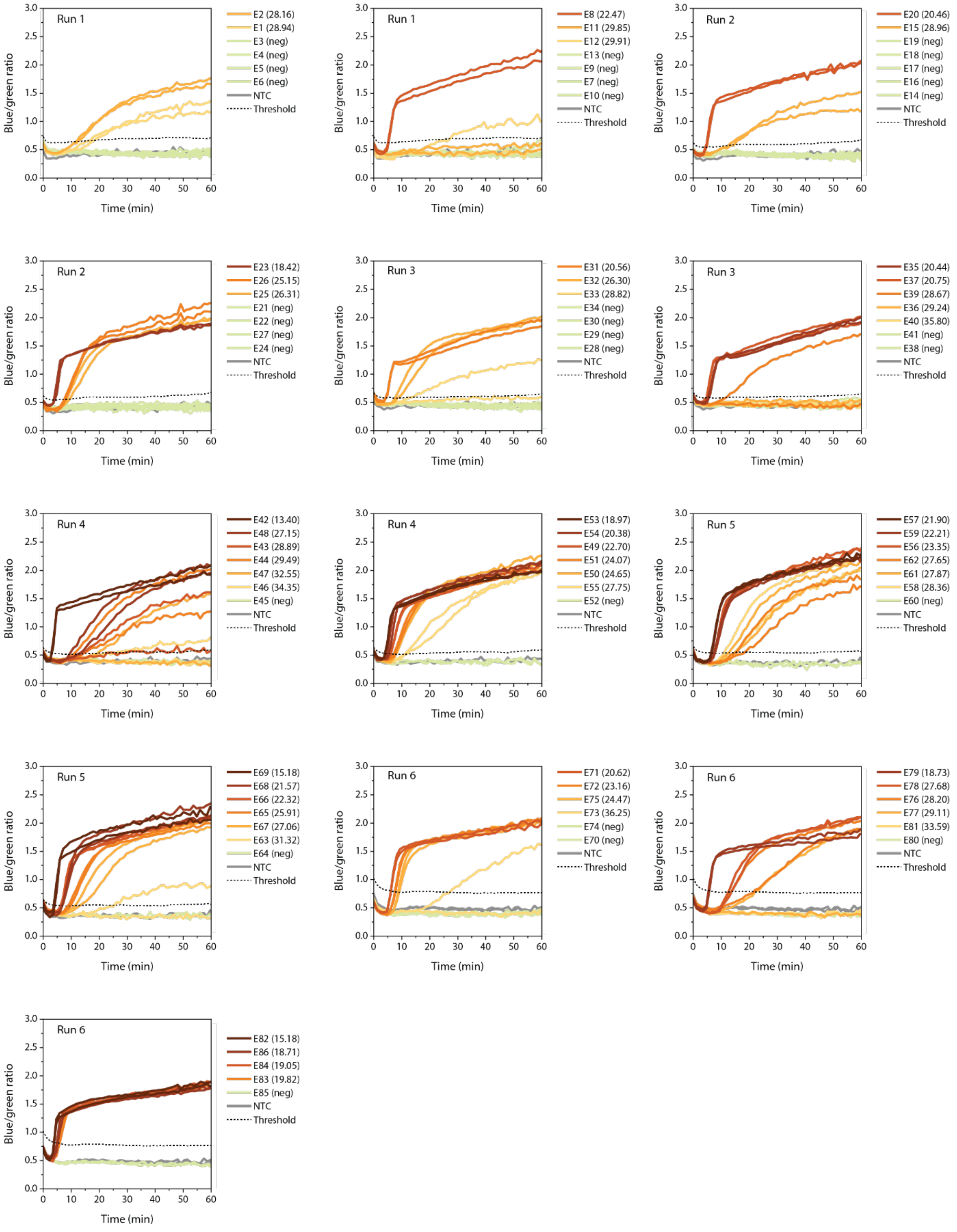

**Supplementary Fig. 12: RT-RPA-LUNAS ratiometric response traces of all extracted clinical samples.** Traces are grouped in panels per assay run, with data divided over multiple panels per run for clarity. In the legend, the sample ID is followed by the corresponding E-gene RT-qPCR $C_t$ value in brackets.

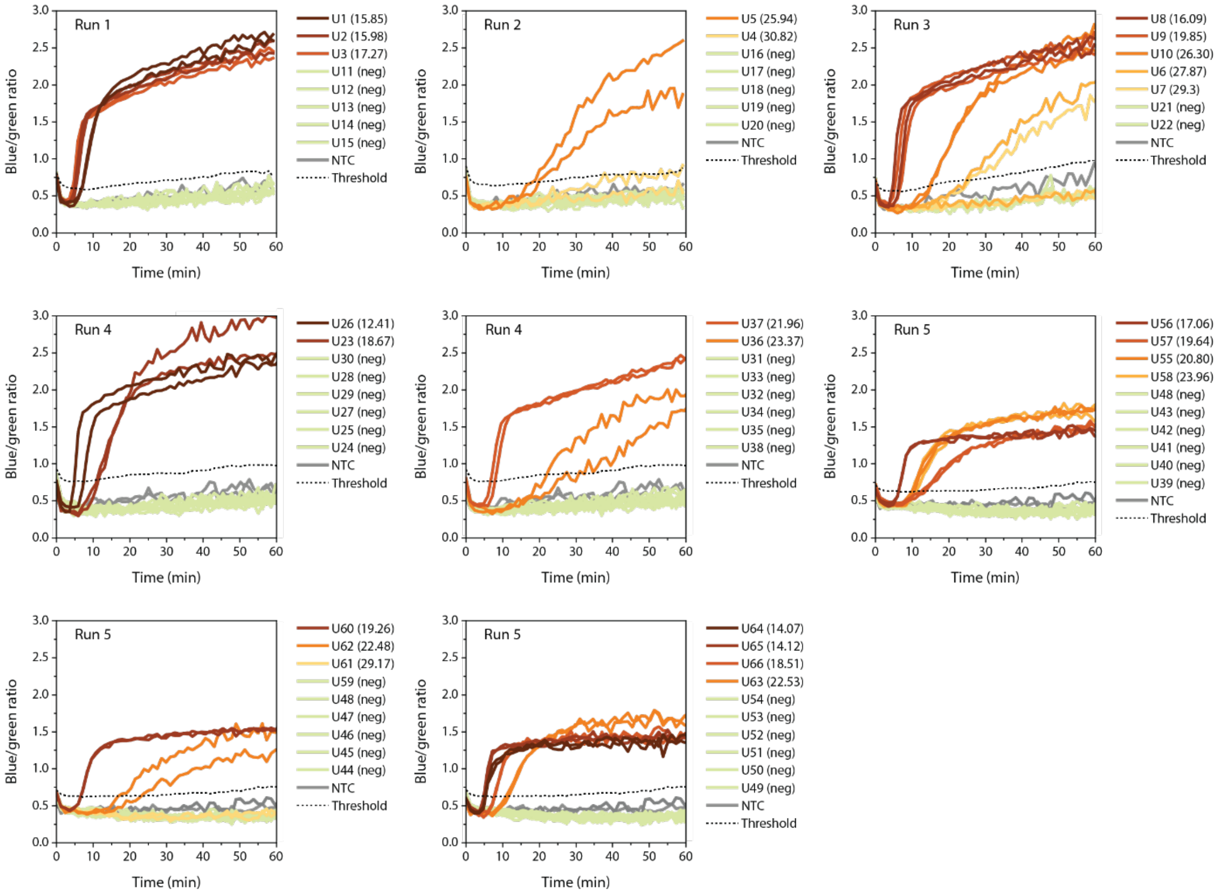

**Supplementary Fig. 13: RT-RPA-LUNAS ratiometric response traces of all unextracted clinical samples**. Traces are grouped in panels per assay run, with data for some runs divided over multiple panels for clarity. In the legend, the sample ID is followed by the corresponding E-gene RT-qPCR $C_t$ value in brackets.

### Supplementary Tables

**Supplementary Table 3: Comparison of RT-RPA-LUNAS SARS-CoV-2 assay performance with that of other recent CRISPR diagnostic methods applied to SARS-CoV-2 detection.**

| Method | Reference | Steps | Readout | Detection time | Approximate LOD |
|---|---|---|---|---|---|
| RT-RPA-LUNAS | This work | 1-pot | Bioluminescent | 10 – 30 min | 200 cp/µL (nasopharyngeal samples, heat-inactivated) |
| SHINE (SHERLOCK based, RPA + Cas13) | Arizti-Sanz et al.[11] | 1-pot | Fluorescent or LFA | 40 min - 1 hour | 100 - 1000 cp/µL (clinical nasopharyngeal samples, heat inactivated)<br>10 cp/µL (synthetic target) |
| SHINE v.2 (SHERLOCK based, RPA + Cas13) | Arizti-Sanz et al.[12] | 1-pot | Fluorescent or LFA | < 90 min. | 200 cp/µL (clinical nasopharyngeal samples, chemically lysed) |
| DETECTR (LAMP + Cas12) | Broughton et al.[13] | 2-step | Fluorescent or LFA | 30 - 40 min | 10 cp/µL (synthetic target), validated for extracted clinical swab samples |
| SCOPE (LAMP + Type III CRISPR (TtCmr) + CARF-RNase) | Steens et al.[14] | 1-pot / 2-step | Fluorescent | 2-step: 35 min<br>1-pot: 180 min | 2-step: 25 cp/µL (synthetic target)<br>1-pot: 482 cp/µL (synthetic target) |
| STOP Covid (SHERLOCK based, RPA + Cas13 | Joung et al.[15] | 1-pot | Fluorescent | 15 – 45 min | 33 cp/mL (with extreme concentration of RNA from clinical samples during magnetic bead isolation) |
| DISCoVER (LAMP + Cas13) | Chandrasekaran et al.[16] | 1-pot | Fluorescent | 60 min | 40 cp/µL (viral stocks in saliva, lysed in integrated device) |
